# Obesogenic diet alters decidual differentiation and cell-cell communication in the mouse uterus

**DOI:** 10.1101/2025.08.13.670199

**Authors:** Burak Koksal, Christian J. Bellisimo, Patrycja A. Jazwiec, Deborah M. Sloboda, Alexander G. Beristain

## Abstract

Maternal obesity is associated with increased risk of infertility, implantation failure, miscarriage, and other pregnancy complications. While prior studies have linked obesity to uterine dysfunction and impaired endometrial biology, how obesity alters the cellular and molecular landscape of the early pregnant endometrium remains poorly understood. Here, we perform single-cell RNA sequencing of embryonic day 5.5 uterus from control and obesogenic mice to generate a cellular atlas of the early decidualizing endometrium. We identify obesity-associated transcriptional changes across multiple Endometrial Stromal Cell (ESC) states and innate immune populations, including uterine natural killer cells and macrophages. Computational modeling reveals that maternal obesity disrupts distinct routes of ESC differentiation during decidualization and leads to shifts in ESC-derived cues known to impact innate immune responses. These findings provide a comprehensive single-cell resource of the post-implantation mouse endometrium while simultaneously generating critical insight into how maternal obesity reprograms the maternal-fetal interface in early pregnancy.

## INTRODUCTION

Early in pregnancy the uterus undergoes dramatic morphological and physiological remodeling to support embryo implantation and fetoplacental development. A central component of this transformation is decidualization, a process where endometrial stromal cells (ESCs) differentiate into specialized decidual stromal cells (DSCs) ^1^. This cellular transformation is accompanied by coordinated changes in the uterine microenvironment, including the recruitment and functional reprogramming of immune cells ^2,3^, extensive vascular remodeling ^4^, and the establishment of a receptive decidual bed ^5^. These processes are tightly regulated and temporally synchronized to facilitate implantation and to promote placental development. In humans, impaired or incomplete decidualization is associated with early pregnancy loss ^6^, implantation failure ^7,8^, and disorders of placentation such as preeclampsia ^9,10^, underscoring its importance not only for pregnancy success but also for the health of the mother and fetus.

Decidual transformation in the mouse is marked by the sequential formation of the primary and secondary decidual zones ^11,12^. Formation of the primary decidual zone (PDZ) precedes the secondary decidual zone (SDZ), initiating between embryonic day (E)4.5-5.0 proximal to the implanted blastocyst at the anti-mesometrial pole. The mature PDZ is composed of tightly packed, non-mitotic decidual cells that form a barrier to immune cell infiltration and protect the early conceptus from oxidative stress ^11^. As pregnancy progresses, a proliferative SDZ develops adjacent to the PDZ from E5.5 onward, expanding toward the mesometrial pole, ultimately contributing to the structural and functional decidua ^11^. These spatially distinct regions play essential roles in modulating the local immune environment, regulating trophoblast invasion, and supporting the conceptus via histotrophic nutrition. While mice and humans differ in the cues initiating decidualization and the spatial distinctions of the PDZ and SDZ, the decidual compartments in the mouse share core physiological functions with the human decidua. Moreover, as in humans, impaired decidualization in mice is a primary driver of local immune dysregulation, placental dysfunction, and adverse reproductive outcomes ^13^.

Differentiation of ESCs into specialized PDZ and SDZ compartments is thought to be orchestrated by a complex interplay of signals derived from multiple maternal and embryonic cell types. These include uterine stromal, immune, epithelial, and endothelial cells, as well as blastocyst-derived trophoblasts, each contributing unique molecular signals that guide differentiation, modulate inflammation, and shape the local uterine environment ^14–16^. While specific factors such as cytokines, growth factors, and extracellular matrix (ECM) components have been implicated in decidual remodeling ^17–19^, our understanding of how diverse cell types coordinate these processes, particularly in the immediate post-implantation period, remains incomplete.

Emerging evidence suggests that metabolic and immune processes profoundly influence decidualization ^20,21^. Metabolic signaling additionally impacts uterine immune cells such as uterine natural killer (uNK) cells and macrophages (Mφ) that play key roles in tissue remodeling and vascular adaptation to pregnancy. Factors that disrupt this relationship can impair decidualization and affect pregnancy success ^22^. In this context, maternal obesity, a chronic, low-grade inflammatory condition, has emerged as a factor that alters tissue metabolism and immune cell function across multiple reproductive compartments ^23–25^. Studies from both human cohorts and animal models indicate that maternal obesity impairs endometrial function, reduces implantation rates, and is linked to higher rates of miscarriage and pregnancy complications such as preeclampsia ^26–29^. While work in IVF clinical populations has underscored the contribution of obesity to endometrial-associated infertility ^30^, the specific cellular and molecular mechanisms by which an obesogenic environment alters key endometrial processes like decidualization during the earliest stages of pregnancy remain poorly understood.

To address this gap, we performed single-cell RNA sequencing (scRNA-seq) of implantation sites collected at E5.5 from control chow and high fat/high sucrose (HFHS) diet-fed mice, a well characterized murine model of maternal obesity^31,32^. This developmental timepoint captures early post-implantation events when decidual zones are forming and extensive remodeling is initiated. This novel dataset offers a comprehensive transcriptional map of uterine cell populations in healthy and obesogenic pregnancy and is available as a web-based resource (*<u>Shiny</u> <u>App</u>*). Our analyses focus on *Hand2*^+^ ESCs, a stromal cell type that initiates decidual transformation, and show that an obesogenic diet disrupts gene signatures central to the control of inflammatory processes, cellular stress responses, and pathways associated with decidualization. Applying single-cell trajectory tools, we show that maternal obesity discretely impacts the development of the PDZ and SDZ. Lastly, we generate a cell-cell interactive model showing how specific cell-cell interactions between endometrial stromal cell types and innate immune cells in the early uterus are potentially altered by maternal obesity.

## RESULTS

### Establishment of E5.5 endometrial single-cell atlases of chow and obesogenic diet exposed mice

To determine how diverse endometrial cell types are affected by maternal exposure to an obesogenic diet, a single-cell transcriptomic approach was applied (Fig. 1A). Briefly, C57BL/6J female mice were subjected to either regular chow (Chow, n = 5) or HFHS (n = 5) diets for 12-15 weeks prior to timed-mating with Chow-fed males. By four weeks of diet exposure, total bodyweight of HFHS mice was greater than Chow controls, and this weight difference was maintained at conception (E0.5) and up to sacrifice at E5.5 (Fig. 1B; Fig. S1A). Along with an overall increase in weight gain, HFHS exposure also led to increases in total body adiposity and peri-gonadal-specific adiposity following ten weeks of diet and at embryonic day (E)5.5 (Fig. 1C; Fig. S1B). The latter increase in gonadal adiposity is consistent with the effects of HFHS diet on metabolic disruption and low-grade inflammation ^33,34^. At E5.5, dams were euthanized and gravid uterine tissues were collected for downstream analysis. Pregnancy was confirmed by the presence of visible uterine implantation swellings and via histology of the paired uterine horn contralateral to that used for single-cell transcriptomics from each dam. Biological characteristics of individual mice are summarized in Table S1.

**Figure 1:**
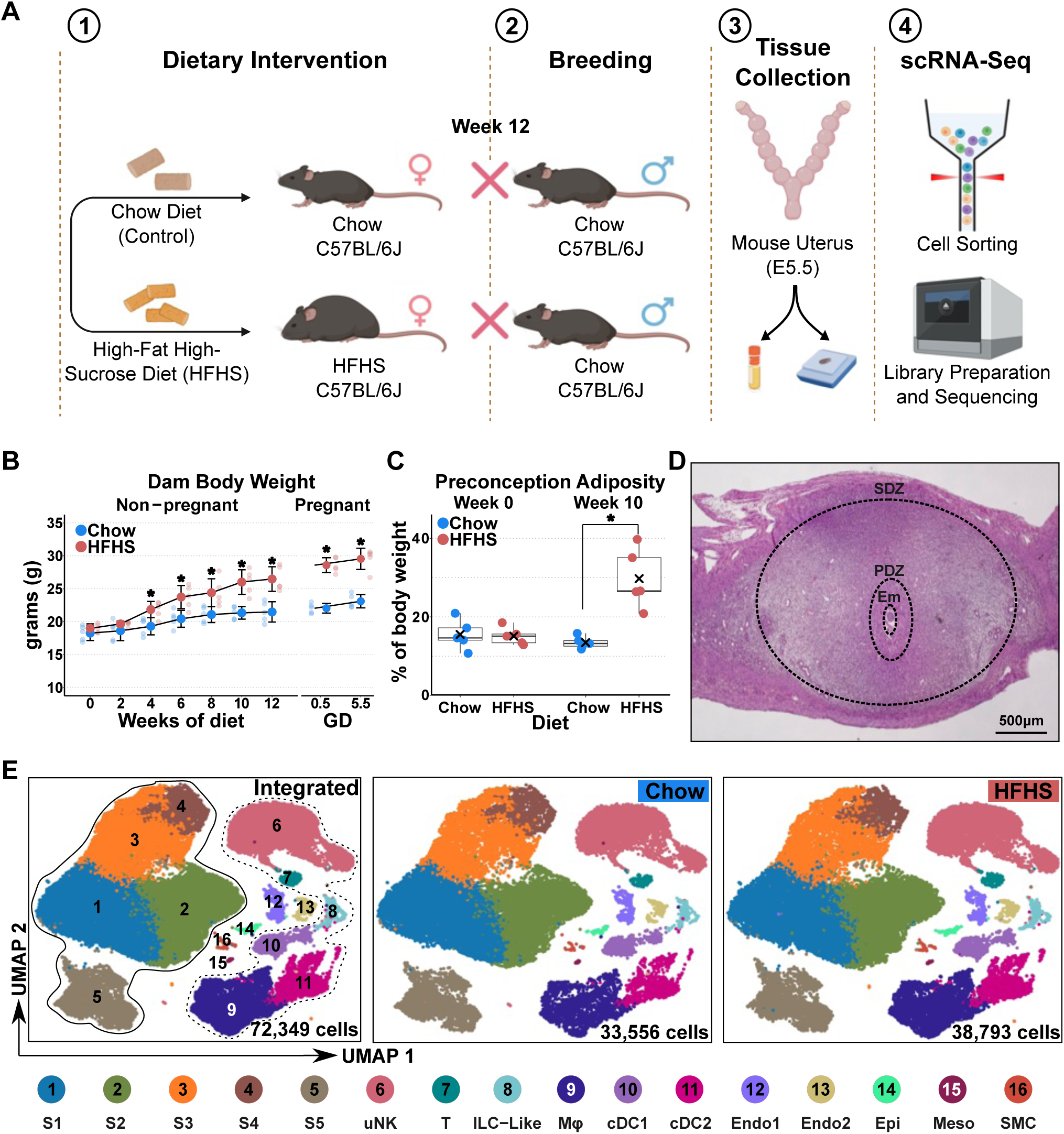
Characterization of obesogenic mouse model and establishment of E5.5 uterine cell atlas (A) Schematic overview of the experimental design. Female C57BL/6J mice were fed a control chow or a high-fat high-sucrose (HFHS) diet for 12 weeks prior to mating (n = 5 per diet group). Uterine horns with intact implantation sites harvested at embryonic day 5.5 (E5.5) were used for histology or cell preparations for single-cell RNA sequencing. (B) Weekly body weight measurements over 12 weeks of dietary intervention. Statistical analyses between groups were performed using repeated measures ANOVA with post-hoc Bonferroni-corrected pairwise comparisons; significance: *p < 0.05. Data shown as mean ± standard deviation; weights of individual dams are depicted. GD = gestational day. (C) Preconception adiposity relative to bodyweight (%) at 0 and 10 weeks of diet. Two-way repeated measures ANOVA was performed; significance: *p < 0.05. (D) Representative brightfield image of an H&E stained E5.5 implantation site; scale bar = 500 μm. Em: embryo; PDZ: Primary Decidual Zone; SDZ: Secondary Decidual Zone. (E) UMAP plot of 72,349 cells from E5.5 implantation sites of Chow (n=5; 33,556 cells) and HFHS (n=5; 38,793 cells) mice. Cell state clusters are indicated as numbers on the UMAP: Endometrial stromal cell clusters (ESC) 1-5 (S1-S5); uterine natural killer cell (uNK); T cell (T); innate lymphoid cell-like (ILC-like); macrophage (Mφ); conventional dendritic cell 1 & 2 (cDC1, cDC2); endothelial cell 1 & 2 (Endo1, Endo2); epithelial cell (Epi); mesothelial cell (Meso), smooth muscle cell (SMC). Solid line outlines clusters specific to ESC states whereas hashed line highlights clusters specific to immune cells.

Single-cell cDNA libraries were prepared and sequenced and following quality control and data pre-processing, 72,349 endometrial cells from the combined Chow (33,556 cells) and HFHS (38,793 cells) groups were integrated and plotted as a Uniform Manifold Approximation and Projection (UMAP) (Fig. 1E). Sequencing metrics showed that percent mitochondrial RNA was not different between individual samples, however, numbers of unique molecular identifiers (UMIs) and number of genes detected did show inter-sample variability, consistent with sample differences in cell capture and sequencing depth (Fig. S1C). Within the combined object, 16 distinct cell clusters were determined, including 6 immune (*Ptprc*^+^) and 10 non-immune (*Ptprc*^-^) cell clusters (Fig. 1E). Leveraging both canonical cell-type markers ^12,35,36^ and the FindAllMarkers function in Seurat ^37^, specific cell types/lineages were identified (Fig. 1E; Fig. S1D). The most prevalent cell types, representing over 70% of all cells captured, were ESCs (S1-5; *Hoxa10*^+^, *Hoxa11*^+^, *Myh11*^-^, *Mcam*^-^), uNK cells (*Eomes*^+^, *Nkg7*^+^), and Mφs (*Adgre1*^+^, *Ms4a7*^+^) (Fig. 1E; Fig. S1D). Additionally, two endothelial cell (Endo-1, −2; *Flt1*^+,^ *CD34*^+^), one epithelial cell (Epi; *Epcam*^+^), one mesothelial cell (Meso; *Upk3b^+^, Lrrn4^+^)*^38^, one smooth muscle cell (SMC; *Acta2*^+^, *Myh11*^+^), two classical dendritic cell (cDC-1, −2; *Zbtb46*^+^), one T cell (T; *Cd3e*^+^), and one innate lymphoid cell-like (ILC-like; *Eomes*^+^, *Cd209a*^+^) types were also identified (Fig. 1E; Fig. S1D; Table S2). Within these subtypes, the Epi cluster contained glandular epithelial (*Foxa2*^+^*, Cxcl15*^+^) and luminal epithelial cells (*Calb1*^+^, *F3*^+^) ^39^, while Endo-1 (*Flt1*^+^, *Mcam*^+^) and Endo-2 (*Prox1*^+^, *Mmrn1*^+^) are likely vascular endothelial ^40^ and lymphatic endothelial cells^35^, respectively (Fig. 1SD). Four ESC clusters (S1-4) expressed *Hand2* transcripts, a marker indicative of ESCs committed to progesterone-induced DSC differentiation (Fig. S1D) ^41^. One *Hand2*^-^ stromal cell population was also captured (S5) and this cell state likely represents a population of deep stromal cells that do not participate in DSC transformation directly but rather play roles in establishing endometrial structural support ^42^. Notably, embryonic (epiblast) and extraembryonic (trophoblast, primitive endoderm) cells of the conceptus were not captured, indicating that the low proportion of this cellular material may preclude sufficient capture following single-cell workflow. Exposure to HFHS diet did not result in significant differences in the proportion of distinct cell clusters or cell types (Fig. S1E), though the proportion of endometrial Mφs trended toward being greater in frequency in HFHS mice (Fig. S1F). Together, we used these data to create a near-comprehensive single-cell resource of the mouse uterus immediately following blastocyst implantation in conditions modeling both healthy weight and maternal obesity and can be explored using a user-friendly web-based interface (*<u>Shiny App</u>*).

### Obesogenic diet drives transcriptomic differences in select E5.5 endometrial cell types

Following the initial characterization of this single-cell dataset, we next set out to examine if an obesogenic diet affected transcriptional readouts in distinct endometrial cell types. Applying a Pearson’s correlation coefficient (PCC) analysis, we first investigated how maternal diet broadly affects endometrial cell gene signatures. This was accomplished by combining all captured cells together followed by comparing the integrated gene signatures of endometrial cells between Chow and HFHS groups. PCC analysis showed that the transcriptome of HFHS endometrial cells was 94.3% similar to that of endometrial cells from Chow fed mice, suggesting that HFHS challenge overall had a modest impact on endometrial cell gene expression (Fig. 2A). Next, a differential gene expression (DGE) analysis was performed comparing the 13 distinct endometrial cell clusters in Chow and HFHS diet groups; for these analyses *Hand2*^+^ S1-4 stromal cell clusters were combined as one cluster. The greatest number of differentially expressed genes (DEGs) were observed in ESCs (*Hand2*^+^ and *Hand2*^-^) and specific immune cell types (i.e., uNK, Mφ, cDC2) (Fig. 2B; Table S3). Specifically, HFHS diet associated with 2063, 308, 307, 219, and 162 DEGs in *Hand2*^+^ stromal, *Hand2*^-^ stromal, uNK, Mφ, and classic DC (cDC2) cells, respectively, with the majority of DEGs showing increased levels in response to the HFHS diet (Fig. 2B). Markedly fewer numbers of DEGs were identified in classic DCs (cDC1; 11 genes), ILC-like (10 genes), T (4 genes), lymphatic endothelial (Endo2; 4 genes), vascular endothelial (Endo1; 3 genes), epithelial (Epi; 1 gene), and smooth muscle (SMC; 1 gene) cells (Fig. 2B). No DEGs were identified in mesothelial cells between diet groups. Importantly, the number of DEGs per cell type/cluster positively correlated with the number of cells per cluster (Fig. S2A), suggesting that in clusters comprised of few cells, the actual impact of diet on gene expression is most likely underestimated. This is a limitation of the single-cell RNA-seq platform and our approach to not enrich for low abundance cell types.

**Figure 2:**
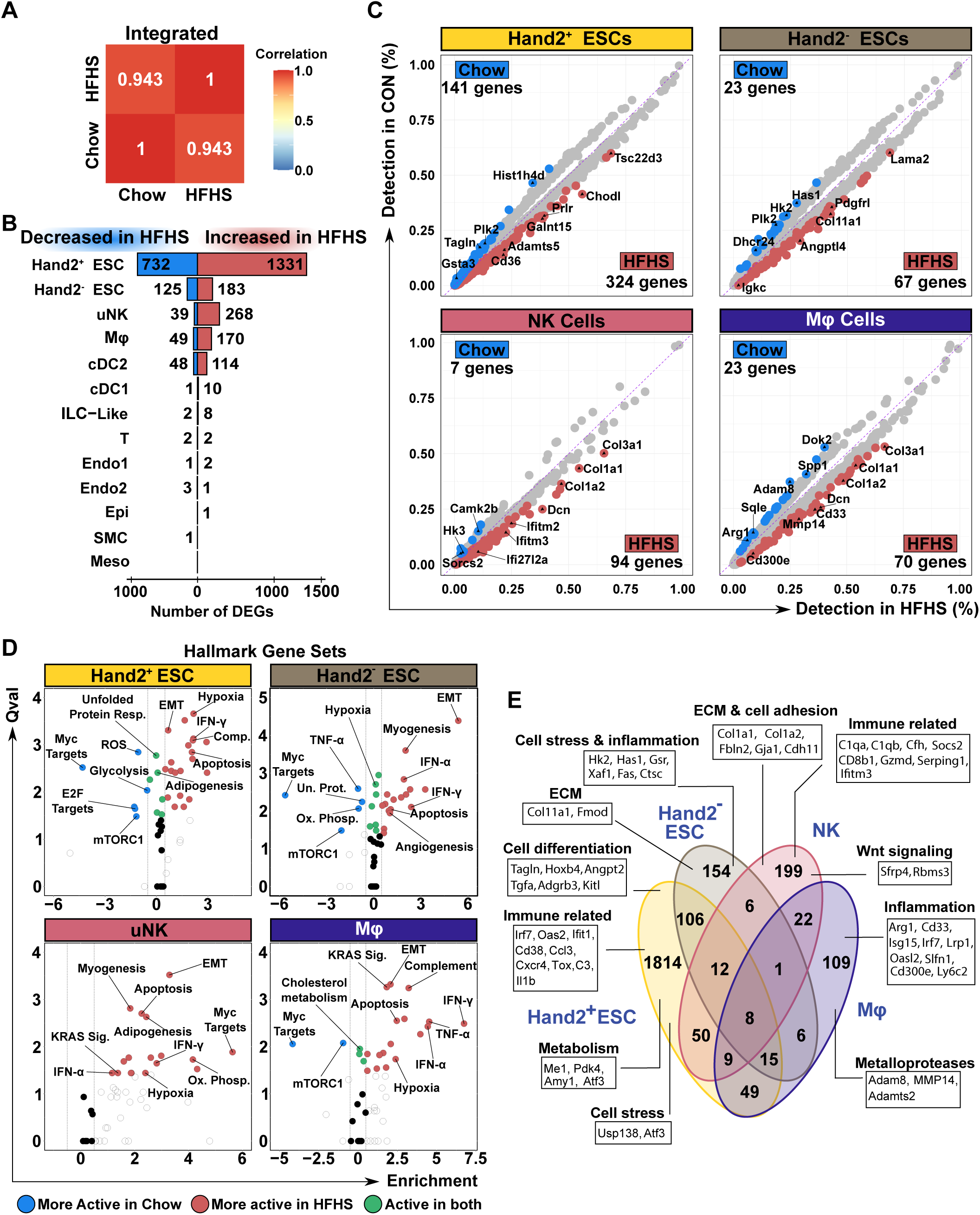
Identification of HFHS-induced transcriptional differences in E5.5 uterine cells (A) Pearson correlation coefficients (PCC) heatmap of Chow and HFHS diet groups. (B) Bar plots showing the number of differentially expressed genes (DEGs) for each uterine cell type as a function of diet. Cess types: Hand2^+^ endometrial stromal cells (ESCs); Hand2^-^ endometrial stromal cells (ESCs); uterine natural killer cells (uNK); macrophage (Mφ); conventional dendritic cell 1 & 2 (cDC1, cDC2); innate lymphoid cell-like (ILC-like); T cells (T); endothelial cell 1 & 2 (Endo1, Endo2); epithelial cell (Epi); mesothelial cell (Meso); smooth muscle cell (SMC). (C) Scatter plots showing DEGs thresholded on adjusted p-val< 0.05 & and absolute Log2FC>0.5 in Hand2⁺ ESCs, Hand2^-^ ESCs, uNK, and Mφ and the detection percentage of each cell type in Chow and HFHS groups. Number of DEGs for each diet group are indicated. (D) Single-Cell Pathway Analysis using Hallmark gene sets obtained from the Molecular Signatures Database for Hand2⁺ ESCs, Hand2^-^ ESCs, uNK, and Mφ. Blue dots signify more activity in Chow, red dots signify enriched activity in HFHS, green dots signify activity in both Chow and HFHS. Qval: Q-value indicating significance; Enrichment indicating extent of pathway enrichment relative to HFHS. Select pathways are highlighted. EMT (Epithelial-to-mesenchymal transition); Ox. Phosp. (Oxidative phosphorylation); ROS (Reactive oxygen species); KRAS Sig. (KRAS signaling). (E) Venn diagram showing DEGs specific to and shared with Hand2⁺ ESCs, Hand2^-^ ESCs, uNK, and Mφ. Examples of genes unique to each cell state are indicated.

In decidualizing *Hand2*^+^ ESCs of HFHS mice, top-ranked DEGs (Adj *P*<0.05; Log_2_FC>±0.4) included genes associated with lipid metabolism (*CD36*), inflammation/immune-related processes (*Galnt15*, *Tsc22d3*) ^20,43^, ECM organization (*Adamts5*) ^44^, and decidual stromal cell functions (*Prlr*) ^45,46^ (Fig. 2C). Genes expressed at lower levels in HFHS *Hand2*^+^ cells associated with cell proliferation (*Plk2, Hist1h4d*) ^47^ and cellular stress (*GSTA3*) ^48^ (Fig. 2C). In *Hand2*^-^ stromal cells (i.e., deep ESCs), genes encoding ECM components show increased levels in HFHS mice (e.g., *Col11a1*, *Lama2*). Additionally, genes associated with vascular development (*Angptl4*, *Pdgfrl*) and pro-inflammatory responses (*Igkc*) are also elevated in HFHS *Hand2*^-^ stromal cells (Fig. 2C). Top genes downregulated in HFHS *Hand2-* stromal cells include genes linked with cell proliferation (*Plk2, Hist1h4d*), ECM turnover (*Has1*), and glucose metabolism (*Hk2*) (Fig. 2C). Like our observations in ESCs, genes showing greatest enrichment in uNK cells and Mφs of HFHS mice included multiple ECM-encoding genes (*Col3a1*, *Dcn*, *Col1a1*, *Col1a2*) (Fig. 2C), indicating the presence of fibrosis-like responses in uterine innate immune cell subsets. Uterine NK cells of HFHS mice showed increases in inflammation-associated genes, with *Ifitm3* and *Ifi27l2a* as top ranked genes (Fig. 2C). Only 3 coding genes showed decreased levels in expression in HFHS uNK cells, and these genes are associated with calcium signaling (*Camk2b*), cell differentiation (*Sorcs2*), and glucose metabolism (*Hk3*) (Fig. 2C). In addition to ECM-associated genes, top genes upregulated in Mφs included the cell membrane-associated metalloprotease *MMP14* and *Cd33,* a gene encoding an immunoglobulin-like receptor important in regulating innate immune cell functions (Fig. 2C).

Single-Cell Pathway Analysis ^49^ showed that HFHS diet resulted in the broad enrichment of pathways central to interferon gamma signaling, hypoxia, cellular apoptosis and epithelial mesenchymal transition (EMT) within multiple cell types (i.e., *Hand2*^+^ stromal, *Hand2*^-^ stromal, uNK, and Mφ) (Fig. 2D; Table S4). Enrichment of the EMT process was most active in *Hand2*^-^ ESCs, while enrichment of the interferon gamma response and other pro-inflammatory responses were top-ranked pathways in uterine Mφs (Fig. S2). Pathways enriched in *Hand2*^+/-^ stromal cells and Mφs of control Chow fed mice included Myc, mTOR signaling, and E2F-directed DNA replication (Fig. 2D), while uNK cells showed specific enrichment of the unfolded protein response pathway (Fig. S2B). Notably, while these endometrium-residing cell types showed enrichment of many of the same gene pathways, the vast majority of DEGs were unique to each cell type with only 8 DEGs shared between *Hand2*^-/+^ stromal cells, uNK cells, and Mφs (Fig. 2E), suggesting that HFHS diet leads to distinct cell-type gene signatures that converge into conserved gene pathways. Together, HFHS-directed transcriptomic differences in E5.5 endometrial cell types suggest that maternal obesity affects multiple cell types, predominately immune and structural/functional stromal cells important in endometrial biology and pregnancy.

### Obesogenic diet disrupts gene pathways central to inflammation and cell stress in *Hand2*^+^ stromal cells

Because the obesogenic diet led to more transcriptomic differences in *Hand2*^+^ stromal cells than in other endometrial cell types, we next set out to more closely examine the effect of HFHS diet on this stromal cell population that is central to DSC transformation. To do this, *Hand2*^+^ cells were subset from the main single-cell object and re-clustered, resulting in five new cell clusters (Fig. 3A). These re-clustered *Hand2*^+^ cells segregated into 1) undifferentiated stromal cells (UnStr; *Vim*^hi^, *Snai2*^hi^, *Igfbp5*^hi^), 2) primary decidual zone cells (PDZ1-2; *H19*^+^, *Dio3*^+^, *Bmp2*^+^), and 3) secondary decidual zone cells (SDZ1-2; *Apoe*^hi^, *Igfbp3*^hi^, *Foxp2*^+^) (Fig. 3A; Fig S3A). The identity of these cell states was determined using previously annotated gene signatures of distinct *Hand2*^+^ cell types ^12,50,51^ (Fig. S3A; Table S5). While the proportions of UnStr cells captured were identical between Chow and HFHS groups, PDZ cell proportions from Chow mice were slightly greater than those found in HFHS mice (∼24% Chow vs 20% HFHS) (Fig. S3B). This contrasted with SDZ cell proportions that showed a moderate increase in HFHS (45%) over Chow (41%) (Fig. S3B). These potential differences in *Hand2*^+^ stromal cell type frequencies suggest that obesogenic diet may affect decidualization in subtle ways impacting both the initial PDZ response as well as the formation of the functional decidua.

**Figure 3:**
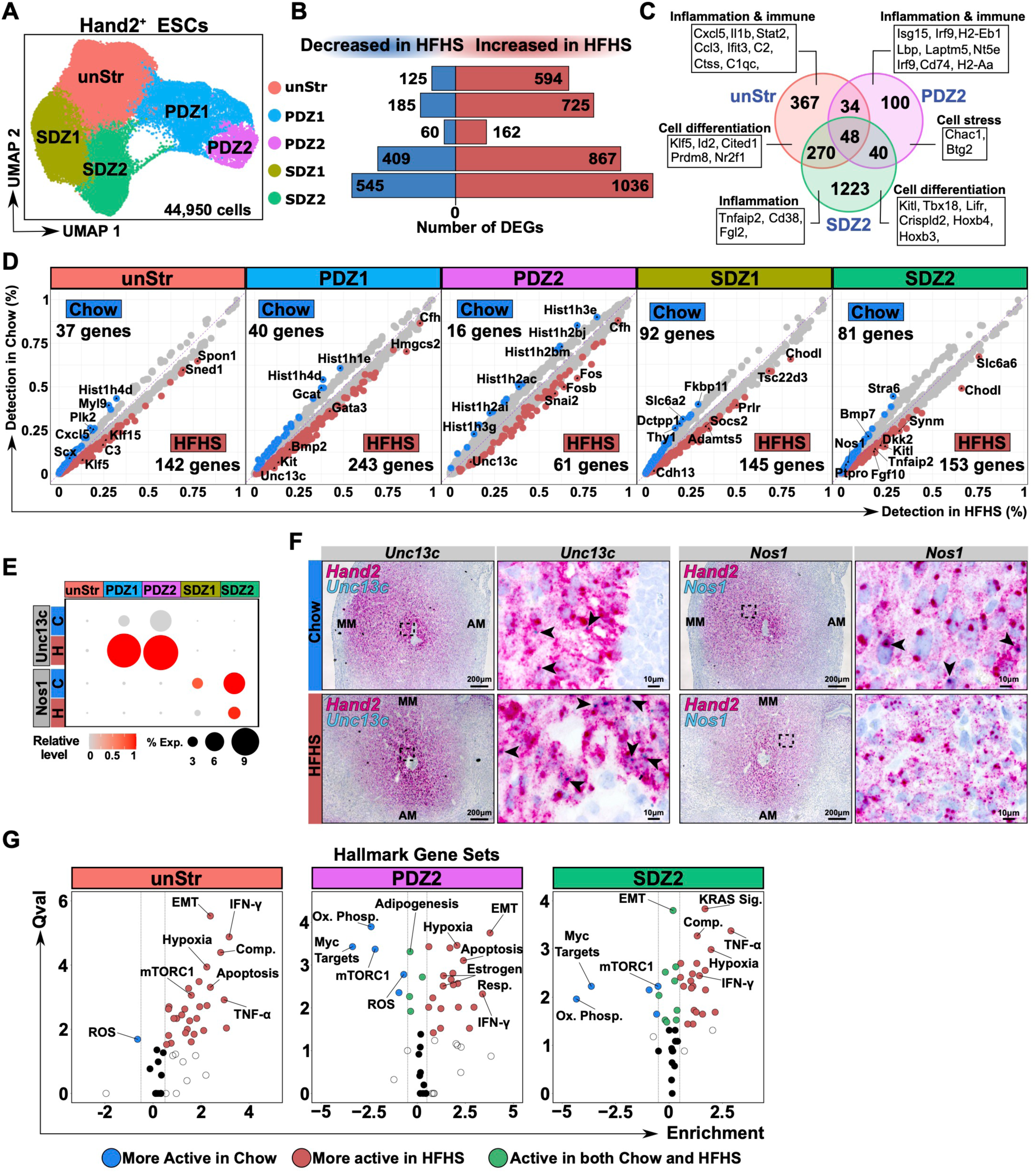
Maternal obesity leads to increased pro-inflammatory and immune-modulating pathways in *Hand2*^+^ ESCs (A) UMAP plot of 44,950 *Hand2*⁺ endometrial stromal cells. UnStr: Undifferentiated Stromal Cells; PDZ1 & PDZ2: Primary Decidual Zone clusters 1 & 2; SDZ1 and SDZ2: Secondary Decidual Zone clusters 1 & 2. (B) Bar plots showing the numbers of DEGs in subtypes of *Hand2*⁺ ESCs. (C) Venn diagram showing unique and shared DEGs expressed in UnStr, PDZ2, and SDZ2 cell states. Highlighted are examples of DEGs specific to each cluster. (D) Scatter plots showing DEGs in *Hand2*⁺ ESC states thresholded on adjusted p-val< 0.05 and absolute Log2FC>0.5. (E) Dot plot showing relative transcript levels of *Unc13c* and *Nos1* in *Hand2*^+^ endometrial stromal cells. The percentage of cluster-specific cells expressing each gene is indicated by dot size. H= HFHS diet; C= Chow diet. (F) *In situ* hybridization images of *Unc13c* (blue signal), *Nos1* (blue signal), and *Hand2* (pink signal) in E5.5 implantation sites. Bar = 100 μm. (G) Single-Cell Pathway Analysis of UnStr, PDZ2, and SDZ2 cell states from Chow and HFHS mice. EMT: Epithelial-to-mesenchymal transition; Comp: Complement; Ox. Phosp.: Oxidative phosphorylation; ROS: Reactive oxygen species; KRAS Sig.: KRAS signaling; Estrogen Resp.: Estrogen response. Blue dots signify more activity in Chow, red dots signify enriched activity in HFHS, green dots signify activity in both Chow and HFHS. Qval: Q-value indicating significance; Enrichment indicating extent of pathway enrichment relative to HFHS.

To examine the effect of HFHS diet on transcriptomic readouts in *Hand2*^+^ cells, we next performed a DGE analysis comparing the five *Hand2*^+^ cell states between Chow and HFHS diet groups. While DEGs were identified in all *Hand2*^+^ cells, the SDZ1 and SDZ2 clusters showed the greatest numbers of DEGs; most genes (SDZ1: 867 of 1276; SDZ2: 1138 of 1581) showing increased levels in response to HFHS challenge (Fig. 3B; Table S6). In UnStr, PDZ2, and SDZ2 stromal cells, 48 DEGs were common to all three cell states, where many of these genes associated with immuno-regulatory functions, including complement activation and antigen presentation (*Cfh*, *Cd74*, *H2-Aa*, *H2-Eb1*, *Ppp3ca*) (Fig. 3C). The majority of DEGs showed cell-state specificity in expression but nonetheless converged into processes common in all three cell types like inflammation and immune function (Fig. 3C). Among top DEGs in UnStr cells (defined as Adj *P*<0.05; Log_2_FC>±0.4), HFHS diet led to increased levels of the complement pathway transcript *C3* and the Krüppel-like transcription factors *Klf5* and *Klf15*, while levels of the chemokine *Cxcl5*, the cell cycle-associated serine/threonine kinase *Plk2*, and the transcription factor *Scx* showed reductions in HFHS (Fig. 3D; Table S6). In the PDZ-1 state in response to HFHS diet, top DEGs showing increased levels include the EMT-regulator *Unc13c* ^52^, the growth factor receptor *Kit*, the transcription factor *Gata3*, and the morphogen *Bmp2*; the latter playing an important role in promoting decidualization ^53^ (Fig. 3D; Table S6). Conversely, top genes showing elevated expression levels in Chow control PDZ1 cells include the glycine acetyltransferase *Gcat*, and nucleosome-associated transcripts *Hist1h1e* and *Hist1h4d* (Fig. 3D). In the PDZ2 state, genes encoding AP1 transcription factors were elevated in HFHS, including *Fos* and *Fosb*, as was the transcription factor *Snai2* and *Unc13c* (Fig. 3D). Like the PDZ1 state, multiple genes encoding nucleosome-associated histone proteins were expressed at higher levels in Chow PDZ2 cells, including *Hist1h2bj*, *Hist1h2ai*, *Hist1h2ac*, *Hist1h3* and *Hist1h3g*, suggesting these cells are actively proliferating or undergoing epigenetic reprogramming (Fig. 3D). In HFHS SDZ1 cells, top genes included *Prlr* (encoding the prolactin receptor), the Jak/Stat pathway inhibitor *Socs2*, the aggrecanase *Adamts5*, and *Chodl,* a transcript encoding a lectin-binding receptor with unknown function ^54^ (Fig. 3D). Top genes enriched in control Chow SDZ1 cells included the cell adhesion molecule *Thy1*, the norepinephrine transporter Slc6a2 and *Dctpp1*, encoding DCTP pyrophosphatase 1 (Fig. 3D). In the SDZ2 cell state, top ranked genes elevated in HFHS included the ligand *Kitl* (encoding stem cell factor), pro-inflammatory *Tnfaip2*, the Wnt inhibitor *Dkk2*, and growth factor *Fgf10*, whereas top genes expressed in Chow SDZ2 cells included the morphogen *Bmp7*, nitric oxide synthase *Nos1*, and *Ptpro* encoding a receptor-tethered protein tyrosine phosphatase (Fig. 3D). *In situ* hybridization (ISH) was performed to validate diet-associated differences in two of the highest ranked DEGs, *Nos1* and *Unc13c in* SDZ and PDZ stromal cell types (Fig. 3E; Fig. 3SB), respectively. ISH confirmed localization of *Nos1* and *Unc13c* to *Hand2*^+^ endometrial stromal cells, with *Nos1* showing specificity to the SDZ and *Unc13c* to the PDZ (Fig. 3F; Fig. S3C). Qualitatively, ISH signals of *Nos1* and *Unc13c* appeared lower and higher, respectively, within endometrial regions of HFHS mice, supporting our scRNA-seq findings (Fig. 3F).

Lastly, Single-Cell Pathway Analysis was performed to further understand the impact of HFHS diet on *Hand2*^+^ ESCs. Like our findings in our single-cell object representing diverse endometrial cell types (Fig. 2E), HFHS challenge led to the enrichment of conserved pathways across all *Hand2*^+^ cell states, including response to hypoxia, the complement cascade, EMT, and IFN-γ signaling (Fig. 3G; Fig S3D; Table S7). In PDZ and SDZ cell types, oxidative phosphorylation, reactive oxygen species, mTor signaling, and Myc signaling pathways were dampened in response to the HFHS diet (Fig. 3G; Fig S3D). HFHS diet also led to the enrichment of other stress- and inflammatory-related pathways in *Hand2*^+^ cells, including TNF signalling, coagulation, UV response, apoptosis, and DNA repair (Table S7), suggesting that exposure to an obesogenic diet establishes an *in utero* environment that imparts conserved inflammatory, cell stress, and metabolic responses in *Hand2*^+^ stromal cell types of the endometrium.

### Obesogenic diet alters PDZ and SDZ differentiation kinetics

To understand how maternal obesity affects the early stages of PDZ and SDZ formation, integrated datasets of Chow and HFHS mice, subset for *Hand2*^+^ ESCs, were subjected to single-cell trajectory analysis. Applying Monocle2 ^55^ pseudotime ordering to *Hand2^+^* cells, a start point enriched in UnStr cells was identified that bifurcated along two distinct pathways culminating in either PDZ2 or SDZ2 cell states (Fig. 4A; Fig. S4A). Along each bifurcated trajectory, *Hand2*^+^ cells progressed first into either PDZ1 or SDZ1 cell states before terminating into either PDZ2 or SDZ2 endpoint states (Fig.4A). Midway through the SDZ trajectory, an additional small branch was observed that showed enrichment of cells expressing senescence-associated genes (Fig. 4A; Fig S4A). These cells are likely senescent stromal fibroblasts and are a byproduct of decidualization ^56^. This novel differentiation path, that PDZ and SDZ cells emerge via distinct routes of differentiation from a common endometrial stromal cell progenitor, is to our knowledge a new observation not previously reported. To more robustly explore this differentiation process, we applied Monocle3 ^57^, another pseudotime ordering tool and showed that similar to Monocle2, UnStr cells localize to the trajectory origin and progress into two major trajectory arms that give rise to either PDZ or SDZ cell states (Fig. S4B). Rendering the Monocle3 trajectory into 3D clearly shows the two distinct arms of *Hand2*^+^ stromal cell differentiation (Figs S4C; S4D). Altogether, these analyses suggest that PDZ and SDZ are developmentally and functionally distinct uterine compartments that can be resolved transcriptionally as early as E5.5 in the mouse endometrium.

**Figure 4:**
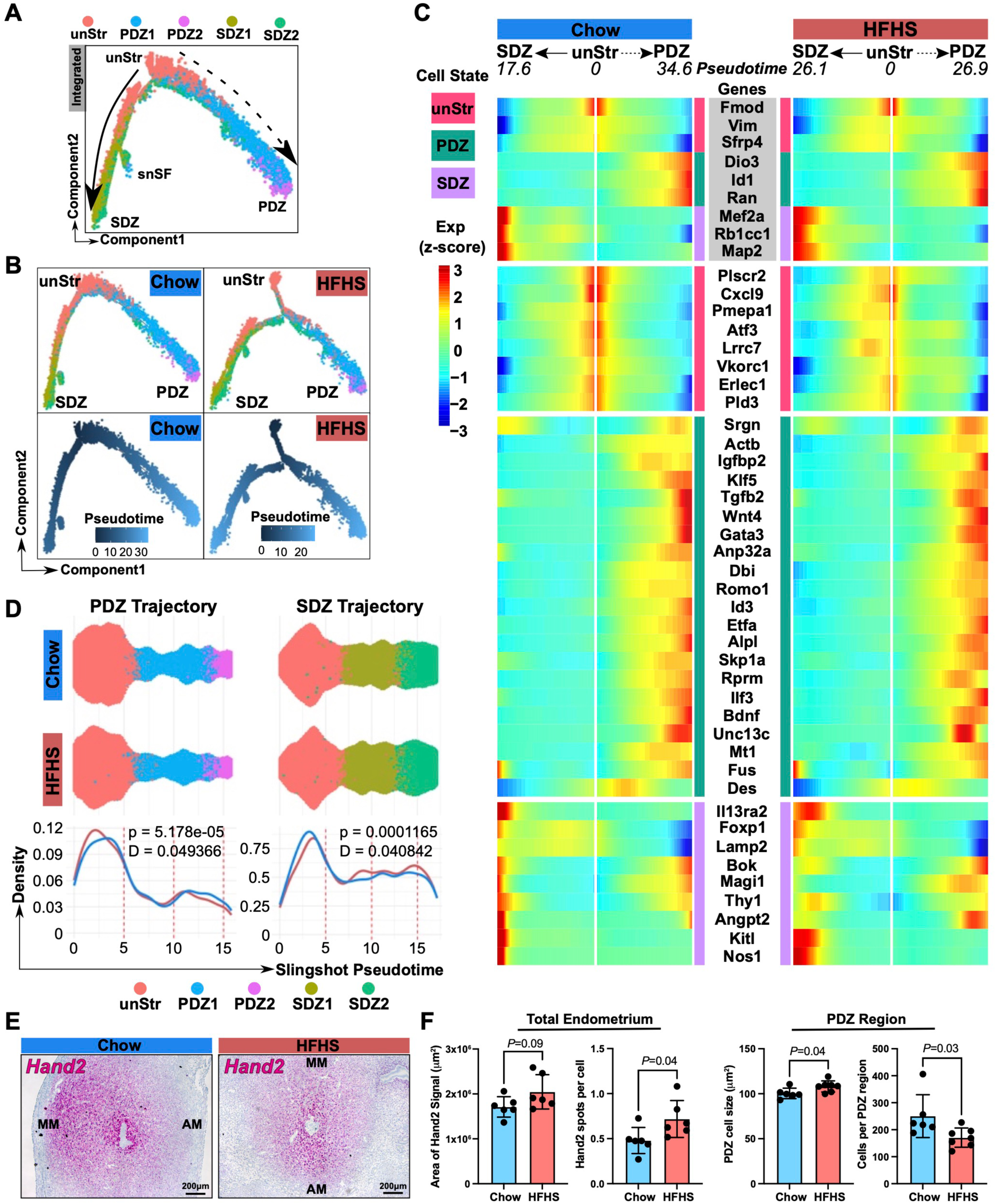
Pseudotime modeling shows altered PDZ and SDZ differentiation kinetics in *Hand2*^+^ cells (A) Monocle2-informed pseudotime ordering of *Hand2*⁺ ESCs from Chow and HFHS mice. UnStr: Undifferentiated Stromal Cells; PDZ1 and PDZ2: Primary Decidual Zone clusters 1 & 2; SDZ1 and SDZ2: Secondary Decidual Zone clusters 1 & 2; snSF: Senescent Stromal Fibroblasts. (B) Monocle2-informed pseudotime ordering of subsetted *Hand2*^+^ cells from Chow and HFHS mice. (C) Heatmaps showing diet-associated patterns of 38 DEGs across Monocle2-inferred pseudotime of *Hand2*^+^ cells from Chow and HFHS mice. A pseudotime value of “0” depicts the origin and arrows indicate direction of cell differentiation from the origin to either the PDZ or SDZ states. Genes used to define pseudotime-predicted differentiation states (UnStr = *Fmod, Vim, Sfrp4;* PDZ = *Dio3, Id1, Ran*; SDZ = *Mef2a, Rb1cc1,* Map2) are indicated with gray background. (D) Swarm and density plots showing cell densities along PDZ and SDZ trajectories constructed using Slingshot. Kolmogorov-Smirnov test was performed to compare distribution between two conditions along the pseudotime. p = *p* value; D = D-statistic. (E) *In situ* hybridization images of *Hand2* transcript in E5.5 implantation sites of Chow and HFHS mice. Bar = 200 μm. MM = mesometrial pole; AM = anti-mesometrial pole. (F) Bar plots showing quantification of *Hand2* signal area (μm^2^) E5.5 implanation sites, number of Hand2 probes per cell in the SDZ region, and cell size/area (μm^2^) and number of cells within the PDZ region. Means +/- standard deviations. Exact p-values are shown.

We then set out to determine whether this bifurcation is vulnerable to maternal diet. Subsetting *Hand2*^+^ cells based on diet and re-applying Monocle2 ordering shows that both Chow and HFHS cells follow a similar bifurcated route of differentiation into PDZ and SDZ cell states (Fig. 4B). However, in the HFHS trajectory, UnStr cells progress along a more elongated path before bifurcating into PDZ and SDZ arms (Fig. 4B). Additionally, the putative senescent stromal fibroblast branch along the SDZ route in HFHS cells occurred earlier in pseudotime in HFHS cells, suggesting that DSC differentiation may be accelerated in this obesogenic model. This interpretation is further supported by the fact that SDZ2 cells of HFHS mice progressed empirically further in pseudotime than SDZ2 cells of Chow mice (Fig. 4B). Correlating the expression of known genes specific to undifferentiated (*Fmod, Vim, Sfrp4)*, PDZ (*Dio3*, *Id1*, *Ran*), and SDZ (*Mef2a, Rb1cc1, Map2*) differentiation states with the pseudotime trajectory confirmed the accuracy of the pseudotime modeling (Fig. 4C). When we correlated *Hand2*^+^ stromal cell trajectories with 38 top-ranking DEGs between Chow and HFHS conditions with branch-specific expression patterns, they showed subtle differences not only in gene level intensity but also at specific points along pseudotime (Fig. 4C). For example, in Chow mice multiple genes (e.g., *Actb, Igfbp2, Romo1*) increased in expression earlier along the PDZ trajectory. This contrasted with the SDZ trajectory, where HFHS cells showed increased levels of most DEGs (e.g., *Il13ra2, Foxp1, Kitl*) earlier along pseudotime. These findings suggest that decidualization kinetics are different in *Hand2*^+^ Chow and HFHS ESCs.

To more thoroughly evaluate how specific *Hand2*^+^ stromal cell states change across pseudotime, we mapped cells onto a Slingshot-inferred trajectory ^58^, which, consistent with our models generated using Monocle2 and Monocle3, identified a bifurcated differentiation pathway originating from UnStr cells and diverging toward either a PDZ or SDZ fate (Fig. S4E). Density-based analysis of cell distributions along pseudotime revealed that at E5.5 the HFHS uterus exhibits a modest but statistically significant enrichment of UnStr cells early in pseudotime and a corresponding depletion of terminally differentiated PDZ cells (Fig. 4D). In contrast, SDZ endpoint cells were slightly more enriched in the HFHS condition (Fig. 4D), suggesting that maternal obesity may bias lineage allocation or alter maturation kinetics across these two discrete decidualization pathways. Our findings suggest that maternal obesity may impair formation of the PDZ and accelerate formation of the SDZ. To assess this latter point, we examined the distribution and signal intensity of *Hand2* transcript, a gene under positive regulation of the progesterone receptor and a marker of decidualization (Fig. 4E). *Hand2* positivity tended to be more profusely distributed in the E5.5 endometrium of HFHS mice, suggesting that decidualization may be increased or accelerated compared to Chow mice, although this difference did not reach statistical significance (Fig. 4E; Fig. 4F). However, the number of *Hand2* transcripts in cells within the SDZ was greater in HFHS mice than in Chow mice, a finding that supports our single-cell modeling suggesting that SDZ development is enhanced in obesogenic conditions (Fig. 4F). Focussing on the PDZ, we next examined the effect of HFHS diet on states of PDZ maturation by measuring cell density and size, since the mature PDZ is composed of densely packed cells arranged around the perimeter of the implantation crypt. In Chow mice, a strong, dense *Hand2* signal was detected in the PDZ while in HFHS mice, *Hand2* positive cells were spatially more diffuse within the PDZ region (Fig. 4E). These qualitative assessments correlated with an increase in cell size and a corresponding decrease in number of cells in the PDZ region in Chow mice compared to HFHS mice, suggesting that PDZ establishment is impaired in obesogenic mice (Fig. 4F). Together our single-cell transcriptomic data and observational histology suggest that maternal obesogenic condition likely modify PDZ development and accelerated the formation of the SDZ at E5.5 in mice.

### Maternal obesity impacts *Hand2*^+^ ESC-directed activities of resident innate immune cells

Our observation that an obesogenic diet alters immune-modulating and inflammation-related gene pathways in ESCs led us to investigate how these changes might influence intercellular communication with key immune cell populations in the E5.5 endometrium. To address this, we applied CellChat analysis ^59^ to our single-cell transcriptomic data to model putative ligand-receptor interactions among relevant cell types. We focused our analysis on *Hand2*^⁺^ ESCs, uNK cells, and Mφs, the three most abundant populations captured in our single-cell dataset and the principal cellular mediators of stromal-immune crosstalk at this developmental stage. We first determined relative changes in both the number and strength of predicted cell-cell interactions across experimental groups (Fig. 5A; 5B; Fig, 5SA). HFHS diet increased the number of signaling interactions directed towards Mφ and SDZ1 cells (Fig. 5A). Conversely, signaling interactions originating from or directed towards PDZ1, PDZ2, and SDZ2 stromal cells were reduced in HFHS implantation sites (Fig. 5A). When evaluating interaction strength, PDZ states in both Chow and HFHS conditions were shown to be the strongest producers of ligands whereas uNK and Mφ immune cell populations showed the greatest receiver potential suggesting the PDZ at E5.5 serves as the primary source of immune-modulating factors (Fig. S5A). Notably, HFHS diet led to marked increases in ESC-directed cell-cell interaction strengths toward uNK and Mφ populations, suggesting an enhancement of immunoregulatory signaling under obesogenic conditions (Fig. 5A). HFHS exposure was predicted to lead to marked increases in uNK-regulating ligands by SDZ2 cells, with high probability of predicted interactions between uNK-expressed nucleolin receptor (*Ncl*) and SDZ2-expressed midkine (*Mdk*) and pleiotrophin (*Ptn*) ligands (Fig. S5B). Additionally, UnStr and PDZ stromal cells from HFHS mice showed marked increases in ligand-directed interactions with Mφs, driven largely by elevated macrophage migration inhibitory factor (*Mif*) and amyloid precursor protein (*App*) ligand expression in UnStr and PDZ cell types with increases in *Cd74*, *Cxcr4*, and *Trem2* expression in Mφs (Fig. S5C).

**Figure 5:**
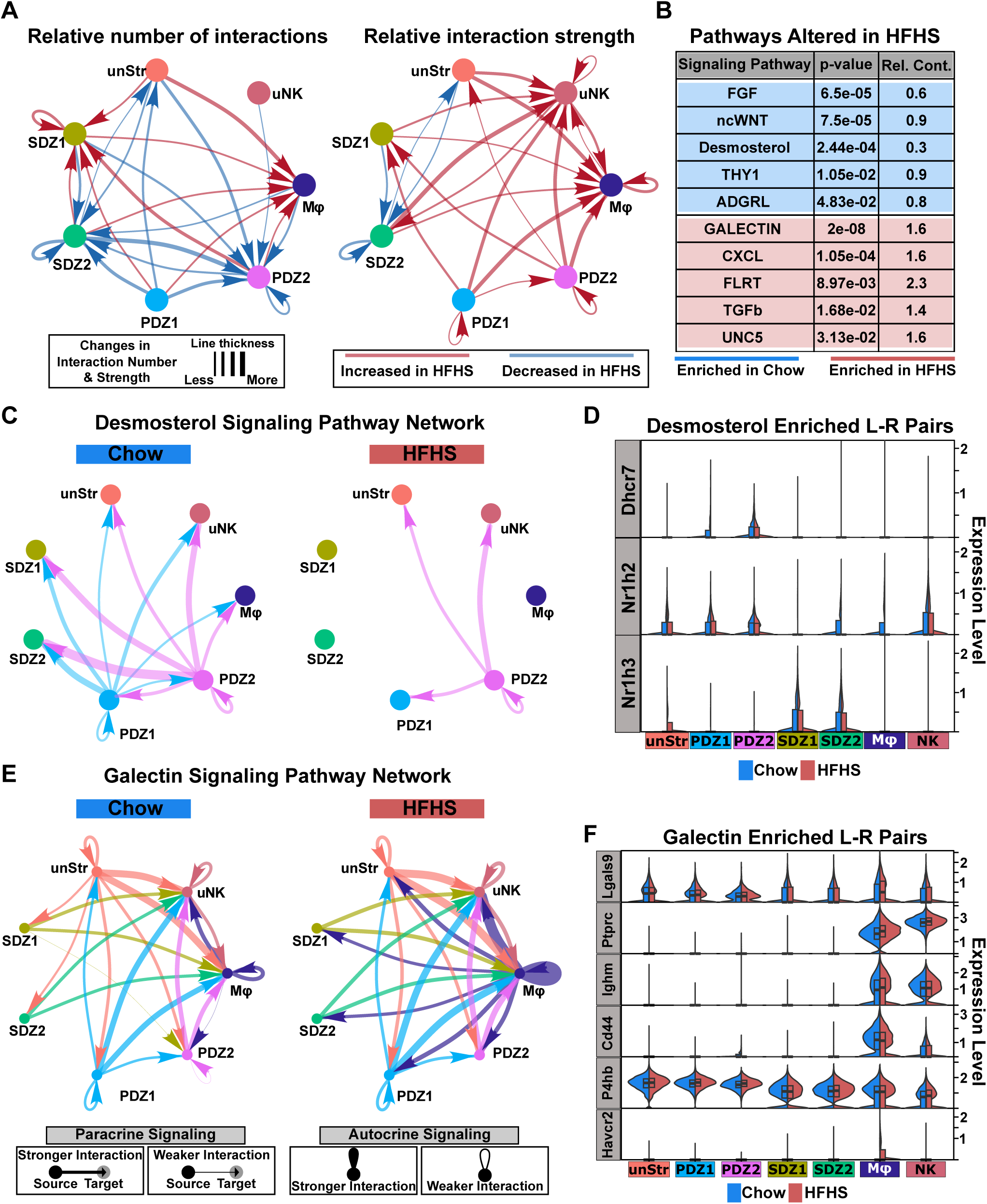
Maternal obesity drives alterations in endometrial stromal cell – immune cell interactions (A) CellChat ligand-receptor analysis of mouse uterine scRNA-seq data. Number and strength of overall interactions between *Hand2*⁺ endometrial stromal cell subsets, uNK, and Mφ. Line colour indicates whether the interaction is increased (red) or decreased (blue) in HFHS. Arrows indicate interaction direction and line thickness correlates the magnitude of interaction number or strength. (B) Top ligand-receptor associated signaling pathways enriched in Chow and HFHS cells. P-value and relative contribution (Rel. Cont.) for each pathway are shown. (C) Circle plots representing the top 50% aggregated interactions for the desmosterol signaling pathway between *Hand2*^+^ cells, uNK, Mφ in Chow and HFHS mice. Line colour indicates the cell type identify of the sender (blue = PDZ1; pink = PDZ2), arrow indicates directionality of the cell-cell interaction, and line thickness correlates with interaction strength. (D) Violin plots showing expression level of ligand/receptor (L-R) DEGs of the desmosterol pathway in *Hand2*⁺ stromal cells, uNK and Mφ. Box plots show 25^th^ and 75^th^ percentiles, and the black line indicates median transcript levels. (E) Circle plots representing the top 50% aggregated interactions for the galectin signaling pathway between *Hand2*^+^ cells, uNK and Mφ in Chow and HFHS mice. Line colour indicates the cell type identify of the sender (red = UnStr; blue = PDZ1; pink = PDZ2; dark green = SDZ1; light green = SDZ2; dark red = uNK; purple = Mφ), arrow indicates directionality of the interaction, and line thickness correlates with interaction strength. (F) Violin plots showing transcript levels of ligand/receptor (L-R) DEGs between Chow and HFHS mice of the galectin signalling pathway. Box plots show 25th and 75th percentiles, and the black line indicates median expression level.

To identify pathways disproportionately altered by HFHS diet, we next ranked signaling pathways based on their relative contributions to overall cell-cell communication under each condition (Fig. 5B; Table S8). Multiple pathways were enriched in HFHS, including the immunomodulatory and inflammation-regulating Galectin and Cxcl-related chemokine pathways and the cell differentiation-associated Fibronectin Leucine-Rich Transmembrane protein (FLRT), TGFβ, and UNC5 (netrin receptor) pathways. In contrast, fibroblast growth factor (FGF), non-canonical (nc)Wnt, and desmosterol signaling were significantly diminished within *Hand2*^+^ stromal, uNK and Mφ cell types (Fig. 5B). Notably, desmosterol signaling, a pathway important in cholesterol biosynthesis and immune cell modulation ^60^, showed a substantial reduction in PDZ cells of HFHS mice (Fig. 5C), suggesting possible diet-dependent suppression of sterol-mediated signaling pathways and desmosterol-directed immune cell dampening in PDZ-stromal communication. To this end, *Dhcr7*, encoding dehydrocholesterol reductase, an enzyme important in cholesterol synthesis, showed decreased levels in HFHS PDZ1 cells, while levels of the transcript encoding Liver X receptor (*Nr1h2*) that binds to and propagates the desmosterol signal, was damped in SDZ2 stromal cells and Mφs (Fig. 5D). In contrast, Galectin signaling, important in driving immunosuppressive signals and shown to impede vascular remodelling in the uterus ^61^, was among the most strongly enriched pathways in HFHS mice, and this was driven primarily by increased paracrine and autocrine-mediated activity from Mφs (Fig. 5E). In these respects, Galectin-9 transcripts (i.e., *Lgals9*) were increased in HFHS Mϕs, while various Galectin-9 receptor transcripts (*Ptprc, Ighm, Cd44, P4hb, Havcr2*) showed increased levels in all *Hand2+* stromal cell types, uNK, and Mφs, the latter pointing to the increase in Galectin autocrine signaling (Fig. 5F).

Together, these findings indicate that HFHS diet exposure reshapes the maternal endometrial communication network during early pregnancy, reducing stromal-derived sterol signaling while enhancing immune-dominated interactions and glycan-mediated pathways between distinct states of *Hand2*^+^ endometrial stromal cells, uNK cells, and decidual Mφs.

## DISCUSSION

Despite growing evidence that maternal obesity impairs endometrial receptivity and decidualization, the cellular mechanisms underlying these effects remain poorly understood. In particular, the impact of obesogenic environments on the post-implantation endometrium has not been examined at single-cell resolution. In this study, we used single-cell RNA sequencing to generate a comprehensive cellular and transcriptomic map of the early decidualizing endometrium in mice fed a control or obesogenic diet. We have also created a publicly accessible web-based resource (*<u>Shiny App</u>*) of the early endometrial cell landscape. We show that maternal HFHS diet leads to enhanced pro-inflammatory and cell stress responses in both ESCs and uterine-resident innate immune cells. Using both computational modeling and *in situ* analyses, we demonstrate that an obesogenic diet leads to changes in the formation of DSCs, as evidenced by altered expression of genes along PDZ and SDZ differentiation trajectories and characterization of PDZ structure. We also show that key cell-cell interactions between *Hand2*⁺ ESCs, uNK cells, and Mφs are disrupted under HFHS conditions, particularly in signaling pathways regulating immune cell activity and tolerance. Together, these findings provide novel insight into how obesity perturbs the cellular architecture and signaling landscape of the uterus during the earliest stages of pregnancy.

Previous studies have consistently reported that maternal obesity is associated with impaired endometrial function, both in natural ^28,62^ and assisted conception contexts ^63^. Work from IVF cohorts shows that obesity negatively impacts endometrial receptivity and embryo implantation rates, independent of oocyte or embryo quality ^27,30^. Transcriptomic analyses of endometrial biopsies from obese women have further identified widespread dysregulation of genes involved in inflammation, oxidative stress, and ECM remodeling ^64^. Complementary studies in rodent models have echoed these findings, with RNA-sequencnig of whole uterine tissue and bulk cell-sorted populations revealing obesity-induced transcriptional shifts in epithelial and stromal compartments, including enrichment of immune-related and metabolic pathways ^65^. However, these prior studies lacked single-cell resolution and were limited to either the pre-implantation period or non-pregnant states, making it difficult to capture pregnancy specific responses like decidualization. Our work addresses this gap by providing the first single-cell atlas of the mouse post-implantation endometrium in the context of maternal obesity.

The PDZ is the first region of the endometrium to undergo stromal cell differentiation following blastocyst attachment, forming a specialized niche critical for protecting the embryo during early pregnancy ^66^. This process is initiated by a convergence of hormonal cues and embryo-derived signals that promote the transformation of ESCs into *Hand2*⁺ DSCs. Notably, we provide evidence using cell trajectory-based inference that PDZ formation is a distinct developmental process to that of SDZ transformation. To our knowledge, distinct DSC differentiation pathways originating from common stromal cell decidual cell precursors has not been previously reported. In light of these novel findings, future work should investigate specific regulators of these disparate developmental pathways as the PDZ and SDZ regions play specialized and distinct roles in embryo development and pregnancy health.

*Hand2* is a key transcriptional regulator of this differentiation process and is expressed in high levels in cells of the early PDZ ^67^. In our study, we used single-cell trajectory analyses to show that the formation of the PDZ is blunted in HFHS mice, where *in situ* analyses examining *Hand2* distribution and expression show that the cellularity and compactness of the PDZ region is altered in HFHS diet-fed mice. These findings suggest that PDZ formation may be impaired or poorly synchronized in the context of maternal obesity. Since our single-cell data revealed that SDZ differentiation may be accelerated in HFHS-fed mice, this raises the possibility that premature emergence of SDZ-like cells disrupts the spatial and temporal fidelity of PDZ formation. However, our histological data suggest this SDZ feature is modest at best. Nonetheless, the consequences of PDZ disorganization could be important as it not only acts as a physical barrier shielding the embryo but also plays a critical immunomodulatory role ^68^, shaping the local uterine immune environment to support embryo tolerance and implantation. If PDZ formation is weakened or spatially dysregulated, ESCs may fail to establish the proper stromal-immune signaling gradients necessary to restrain inflammatory immune activity in the immediate peri-embryonic space. This possibility is supported by our identification of altered expression of immune-regulatory transcripts such as multiple AP1 transcription factor-encoding transcripts and multiple interferon-response genes (ISGs) within the decidualizing stroma, as well as disrupted ligand-receptor interactions between *Hand2*⁺ PDZ cells, uNK cells, and decidual macrophages in HFHS mice. Together, these findings suggest that maternal obesity perturbs early decidual organization in a way that may compromise the immunological integrity of the maternal–fetal interface. In addition to revealing shifts in stromal differentiation dynamics, our single-cell communication analyses identified diet-induced disruptions in specific paracrine signaling pathways between ESCs and uterine immune cells. Notably, CellChat analyses showed that desmosterol signaling, originating primarily from PDZ stromal subsets, was markedly reduced in HFHS-exposed mice. Desmosterol, a cholesterol biosynthesis intermediate, acts as a ligand for liver X receptors (LXRs), which regulate inflammatory responses in macrophages and other immune cells ^60^. Its reduction may therefore contribute to a pro-inflammatory shift in the local decidual environment. In contrast, Galectin signaling, particularly through Galectin-9 (*Lgals9*), was enriched under HFHS conditions. Galectin-9 is a glycan-binding protein with complex immunoregulatory roles, including modulation of T cell exhaustion, NK cell cytotoxicity, and Mφ polarization ^69,70^. Enhanced Galectin-9 signaling from decidualizing ESCs toward uNK and macrophage populations may reflect a compensatory or dysregulated attempt to restrain inflammation in the context of maternal metabolic stress. Together, these HFHS-induced ligand-receptor differences highlight how maternal obesity reshapes the immune-regulatory signaling landscape of the decidua, potentially altering the balance between pro- and anti-inflammatory cues in its early formation.

While maternal obesity is a well-established risk factor for adverse pregnancy outcomes in humans, our HFHS mouse model does not recapitulate all aspects of this clinical phenotype. Although HFHS-fed females exhibit reduced fertility ^71^, requiring more mating attempts to establish pregnancy, the number of implantation sites and fetal and placental weights at E17.5 are not significantly different from Chow-fed controls ^72^. These observations are consistent with some, but not all reports in the literature, where variability in diet composition, strain background, and environmental factors likely contribute to divergent outcomes. We deliberately chose a standard chow diet as a control, rather than a purified, calorie-matched control diet, because this better recapitulates the human context. While nutrient-matched control diets are optimal for isolating the effects of specific macronutrients, they do not reflect the broader dietary shifts that occur in human populations consuming high-fat, energy-dense foods that results in increased adiposity. In humans, high-fat diets are rarely consumed in isolation but are accompanied by changes in micronutrient density, fiber type, and food processing. By comparing a high-fat diet to standard chow, we model the combined nutritional changes typical of real-world diets, prioritizing translational relevance over strict nutrient matching. Despite the absence of gross pregnancy defects, our study reveals substantial obesity-related alterations in differentiation dynamics and intercellular signaling within the peri-implantation endometrium. These findings raise the possibility that maternal obesity may induce subtle or latent disruptions in uterine biology that are not sufficient to compromise fetal growth under controlled laboratory conditions but could render pregnancies more vulnerable to additional environmental or physiological stressors. Indeed, our prior work has shown that maternal HFHS exposure impairs spiral artery remodeling at mid-gestation (E10.5), although these defects normalize by E17.5 ^73^. Together, these findings suggest that the effects of maternal obesity may be developmentally transient or context-dependent, with compensatory mechanisms buffering the uterine environment in the absence of secondary insults. Future studies incorporating additional stressors, such as infection, social stress, or hypoxia, may help reveal the full extent to which an obesogenic uterine environment sensitizes pregnancies to failure or dysfunction, providing a more faithful parallel to the complex and multifactorial nature of human pregnancy complications.

## Limitations of the study

This study uses a mouse model of diet-induced obesity, which, while valuable for dissecting mechanistic pathways, does not fully recapitulate the complexity of human obesity, including its metabolic, hormonal, and inflammatory diversity. Additionally, our use of single-cell RNA sequencing is limited by biases in cell capture and detection sensitivity. Rare and fragile cell populations may be underrepresented, affecting both cluster resolution and statistical power in differential expression analyses. Moreover, single-cell transcriptomics lacks spatial context, which is a notable limitation in the uterus—a highly structured and spatially dynamic tissue.

## ACKNOWLEDGEMENTS

We thank Dr. M Shannon for his help with *in situ* hybridization. We thank members of the Beristain laboratory, specifically Matthew Shannon, Gina McNeill, and Victoria Leonard, for their critical reading and feedback of this manuscript. We additionally thank the computational resources provided by the Advanced Research Computing Team, University of British Columbia.

## AUTHOR CONTRIBUTIONS

Conceptualization: A.G.B., C.J.B., D.M.S.; Methodology: B.K., C.J.B., P.A.J., D.M.S., A.G.B.; Formal analysis: bioinformatic – B.K., animal work and benchwork – B.K., P.A.J., C.J.B.; Resources: A.G.B., D.M.S.; Writing: original draft – B.K., A.G.B.; review & editing – B.K., C.J.B., D.M.S., A.G.B.; Supervision: A.G.B., D.M.S.; project administration: A.G.B. Funding acquisition: A.G.B., D.M.S.

## SUPPLEMENTAL FIGURE TITLES AND LEGENDS

**Figure S1:**
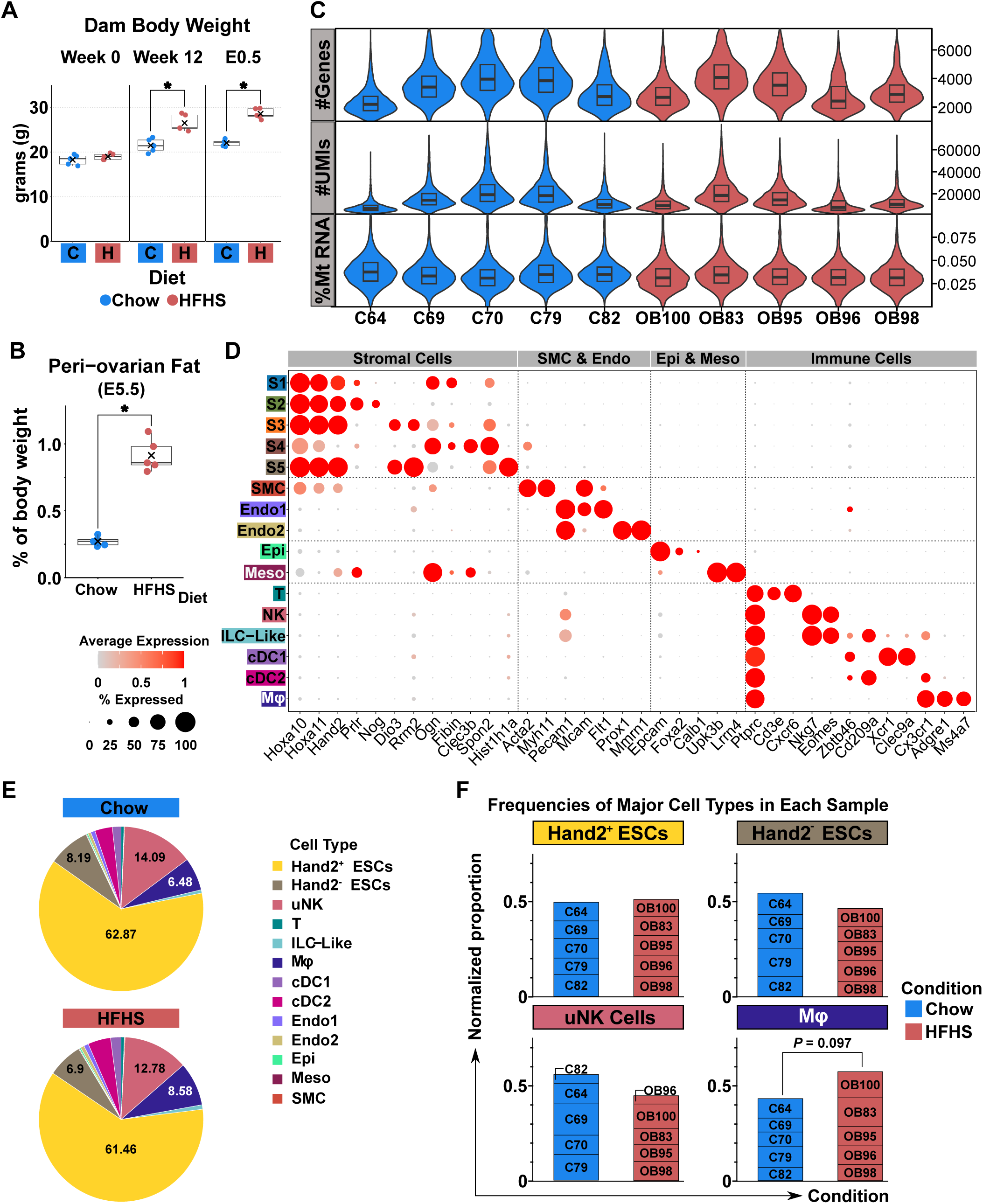
Obesogenic mouse model assessment and single-cell data quality control (A) Body weight measurements in grams (g) of dams subjected to 0 and 12 weeks of diet pre-pregnancy and at plug detection (E0.5). Statistical analyses between groups were performed using two-way ANOVA followed by Tukey’s multiple comparisons test. *= p < 0.01. (B) Peri-ovarian adiposity as percent of total body weight following 12 weeks Chow or HFHS diet at E5.5. Analyses between groups were performed using Welch’s t-test. * = p<0.0001. (C) Violin plots showing the distribution of the number of genes (nFeature_RNA), Unique Molecular Identifiers (#UMIs, nCount_RNA), and mitochondrial DNA percentage (%MtRNA) after quality control and data pre-processing for each sample. Box plots indicate 25th and 75th percentiles, and the black line indicates data median. Individual samples are denoted on the x axis (C = Chow or OB = obesogenic diet). (D) Dot plot showing the average log expression level (colour) and proportion of cells expressing (dot size) of select genes used for determining the cell type. (E) Pie charts showing the proportion of each cell cluster in Chow and HFHS groups. (F) Stacked bar plots plotting the frequencies of *Hand2*^+^, *Hand2*^-^, uNK, and Mϕ cell types in Chow and HFHS mice. Data analyzed using Welch’s t-test.

**Figure S2:**
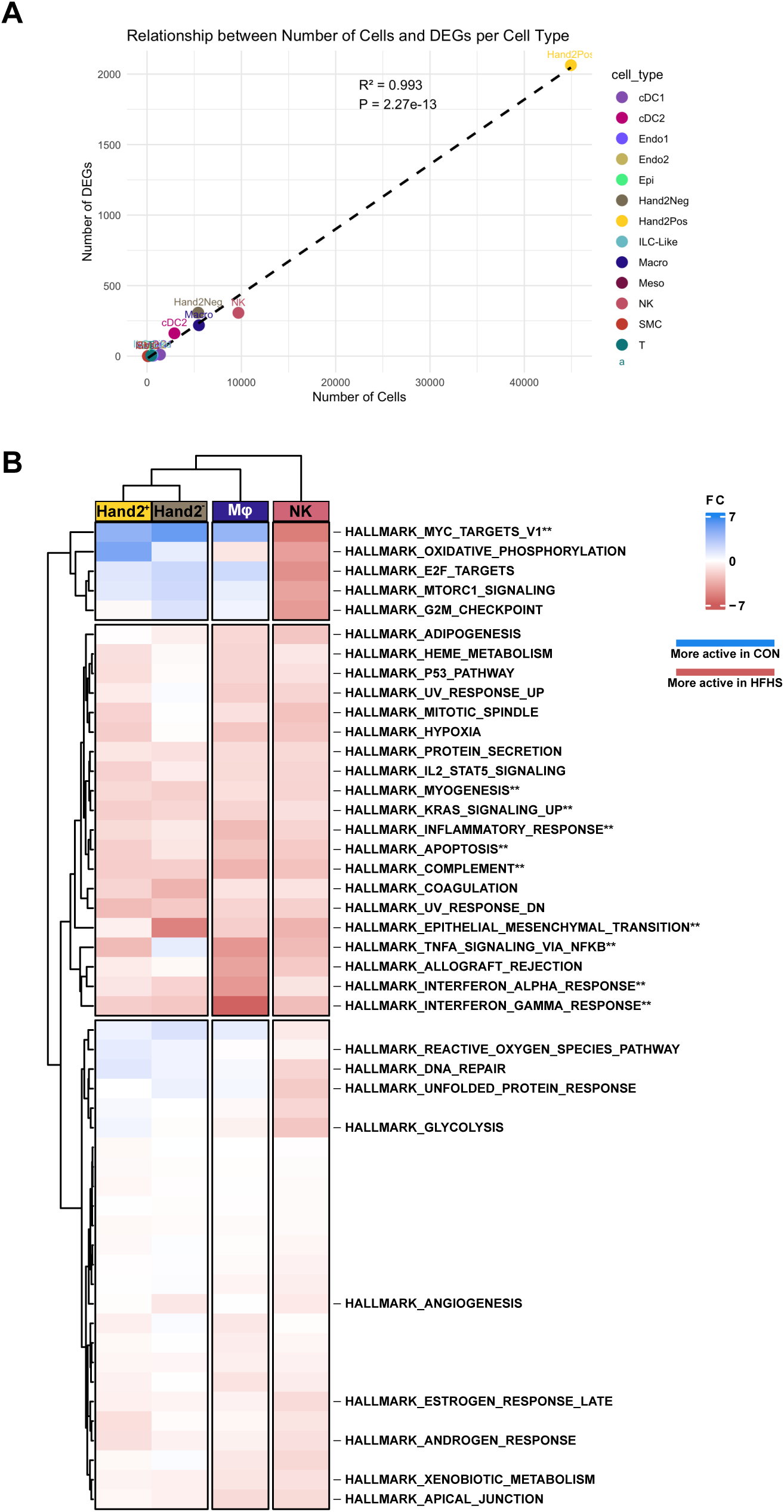
HFHS-induced changes in gene pathways of *Hand2*^+/-^ stromal and uNK and Mθ> cell types (A) Linear regression showing the linear relationship between number of captured/sequenced single cells and the number of differentially expressed genes (DEGs) in cell type/clusters determined in single-cell transcriptomic analysis. Shown is the coefficient of determination (R^2^) and P-value. (B) Heatmap showing absolute fold-change differences of Hallmark gene sets showing enrichment in *Hand2*^+^, *Hand2*^-^, uNK, and Mϕ cell types in relation to Chow or HFHS diet. Blue = more active in Chow; Red = more active in HFHS. ** denotes pathways significantly different (adjusted p-value < 0.01; absolute FC > 0.5) between Chow and HFHS diet conditions for each cell type.

**Figure S3:**
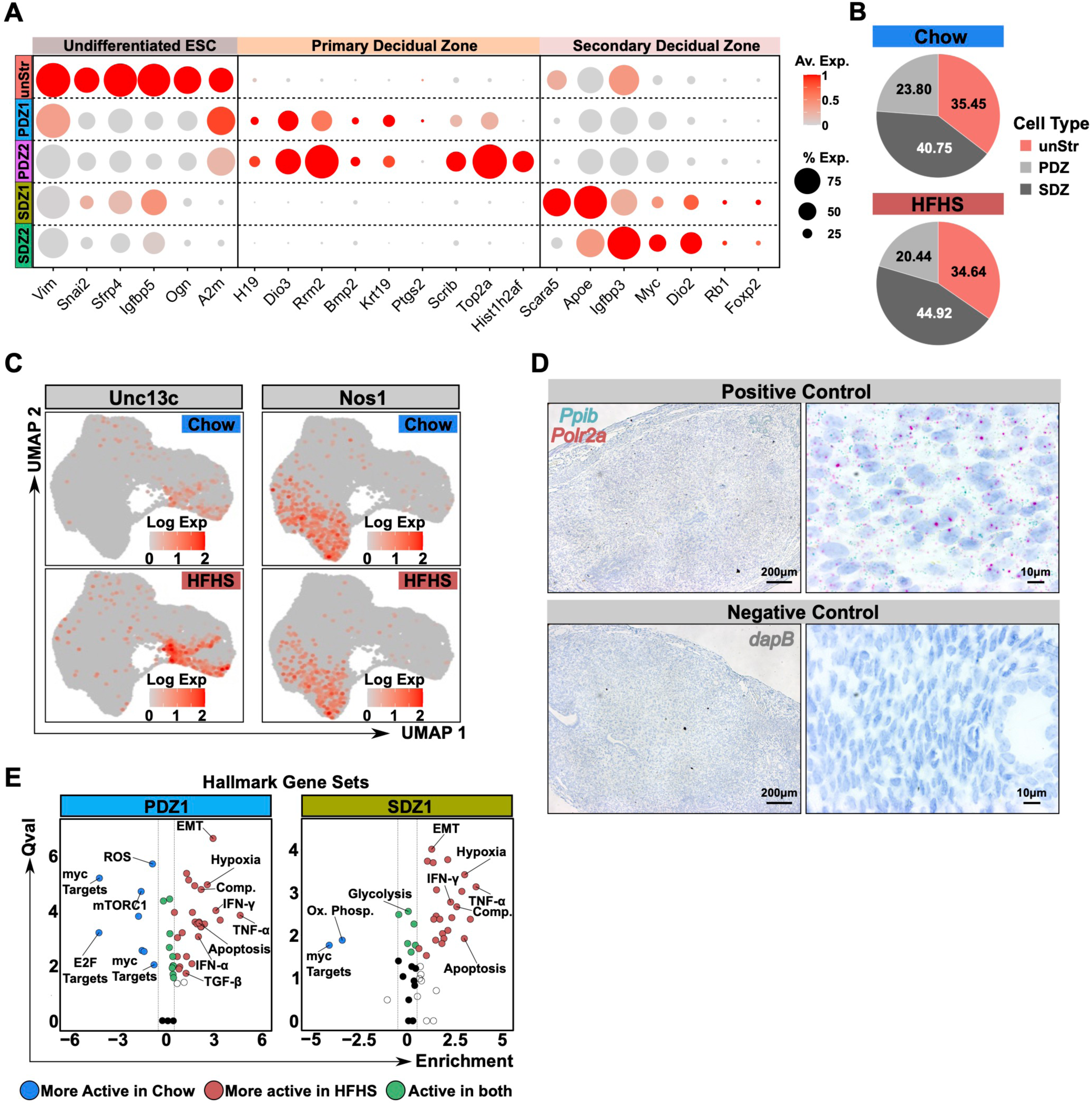
Cell subset identification, pathway analyses, and gene verification in Hand2⁺ stromal cells (A) Dot plot showing average log-transformed transcript levels (Av. Exp) of select genes used in identifying cell identities of *Hand2*^+^ undifferentiated endometrial stromal cells (ESC), primary decidual zone stromal cells, and secondary decidual zone stromal cells. The proportion of cells (% Exp) within each cell state expressing each gene is denoted by dot size. (B) Pie chart depicting the proportions of undifferentiated endometrial stromal cells (UnStr), primary decidual zone cells (PDZ), and secondary decidual zone cells (SDZ) in Chow and HFHS mice. (C) UMAP plots showing log-transformed mRNA levels of *Unc13c* and *Nos1* in Chow and HFHS in *Hand2*^⁺^ endometrial stromal cell subsets in UMAP space. (D) *In situ* hybridization images of positive (*Ppib*, blue signal; *Polr2a,* red signal) and negative (scrambled oligo) control probes in E5.5 implantation sites. Bar = 100 μm. (E) Single-Cell Pathway Analysis showing enriched Hallmark gene sets in PDZ1 and SDZ1 *Hand2*^+^ cell states (Adjusted p-value < 0.01, absolute FC > 0.5). Blue dots = more active in Chow; Red dots = more active in HFHS; Green dots = active in both Chow and HFHS. Qval: Q-value indicating significance level. Select pathways are highlighted by lines. EMT: Epithelial-to-mesenchymal transition; Comp: Complement; Ox. Phosp.: Oxidative phosphorylation; ROS: Reactive oxygen species pathway.

**Figure S4:**
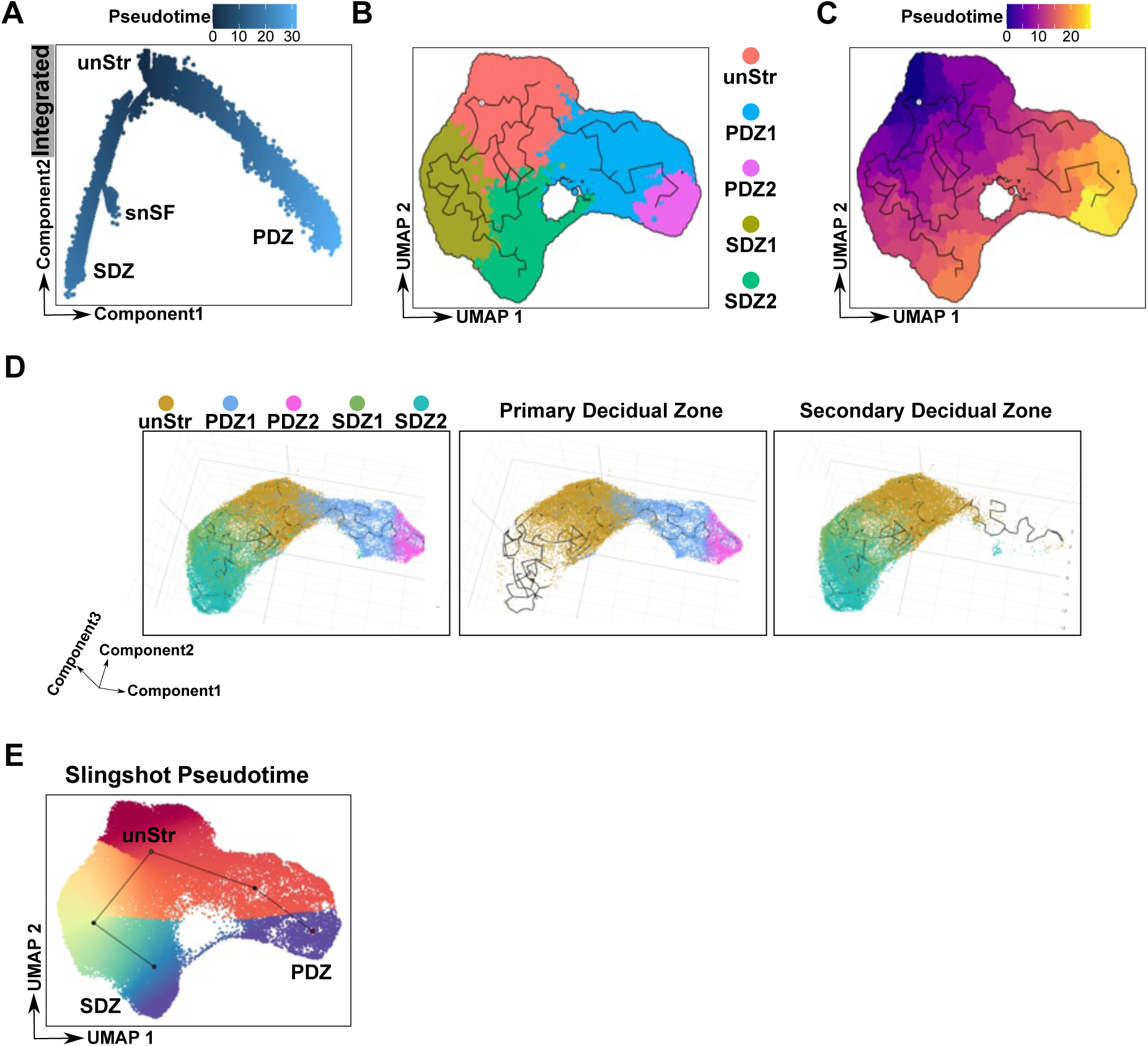
Monocle2, Monocle3, and Slingshot pseudotime ordering of *Hand2*^+^ stromal cells (A) Monocle2 pseudotime ordering of Chow and HFHS (integrated) *Hand2*⁺ stromal cells. Shade of cell along the trajectory denotes pseudotime value. UnStr: Undifferentiated Stromal Cells; PDZ1 & PDZ2: Primary Decidual Zone clusters 1 & 2; SDZ1 & SDZ2: Secondary Decidual Zone clusters 1 & 2, snSF: Senescent Stromal Fibroblasts. (B) UMAP of *Hand2*^+^ stromal cells overlain with Monocle3-determined cell trajectories (black lines). (C) Monocle3-determined pseudotime heatmap color gradient showing pseudotime values within UMAP space. (D) 3-Dimensional rendering of *Hand2*⁺ stromal cell UMAP overlain with Monocle3-informed lineage trajectories (black lines) showing PDZ and SDZ branches together or separately. (E) UMAP of *Hand2*⁺ stromal cells overlain with Slingshot-determined lineage trajectories (black lines). Cells are colour coordinated by estimated Slingshot pseudotime (red = early; blue = late) for integrated object.

**Figure S5:**
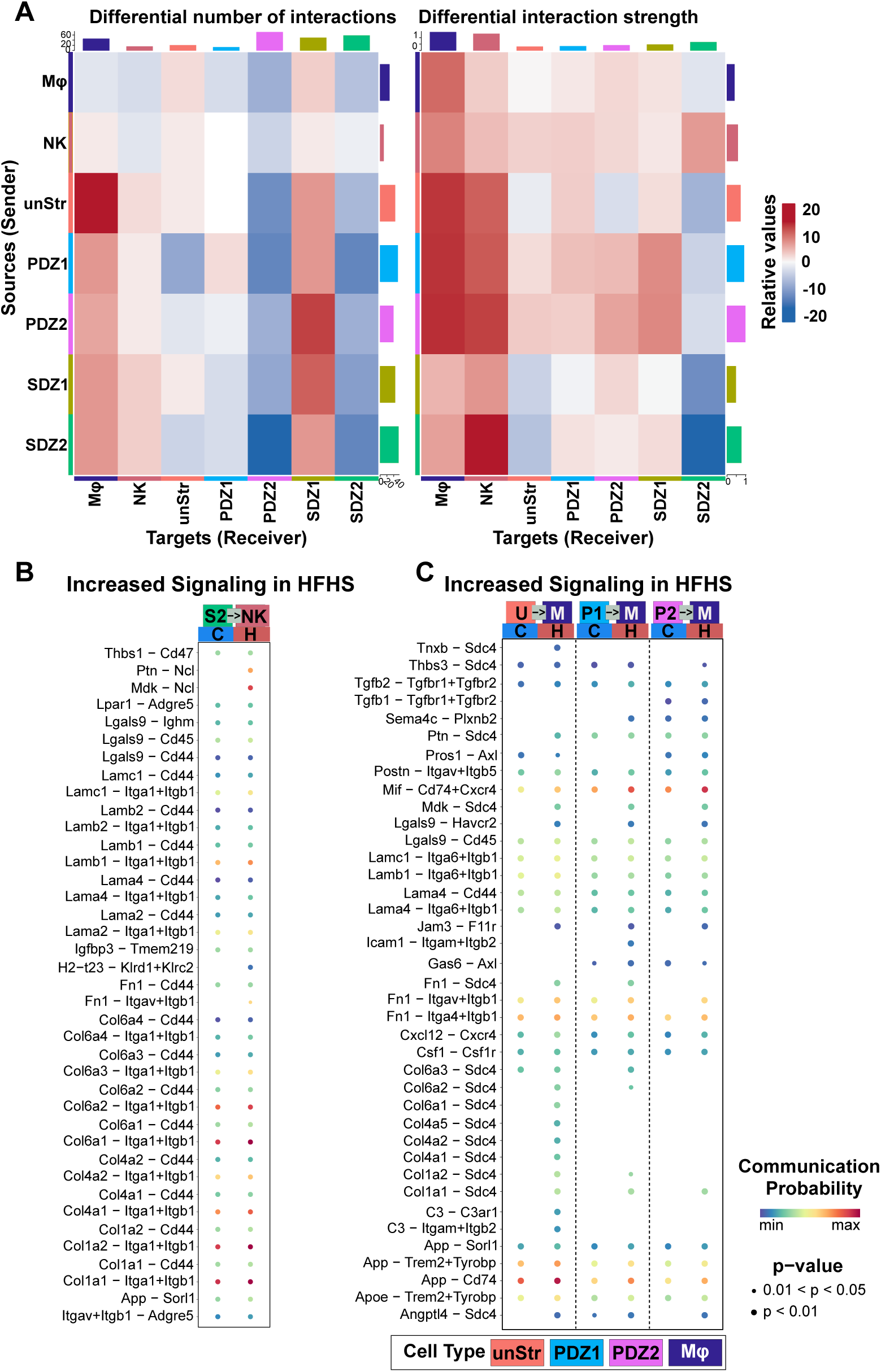
Identification of diet-driven alterations in cell-cell ligand-receptor interactions (A) Heatmap shows the net change in number of predicted ligand-receptor interactions (CellChat) between sender (y-axis) and receiver (x-axis) cell populations, comparing HFHS to control conditions. Red and blue represent gains and losses of interactions under HFHS, respectively. Bar graphs at the margins indicate the total number of increased or decreased interactions per cell group, with values representing the total number of differentially gained or lost ligand-receptor interactions. Significant ligand (L)-receptor (R) pairs between (B) SDZ2 (S2) and uNK (NK) and (C) UnStr (U), PDZ1 (P1), PDZ2 (P2), and Mφ (M) cell types contributing to increased signaling in HFHS diet. Dot color indicates communication probabilities (blue = low; red = high) and dot size indicates the p-value (one-sided permutation test) of the activity of the L-R driven pathway in Chow or HFHS conditions. Arrows indicate directionality of the ligand - receptor interaction between cell types.

## METHODS

### Key Resources Table

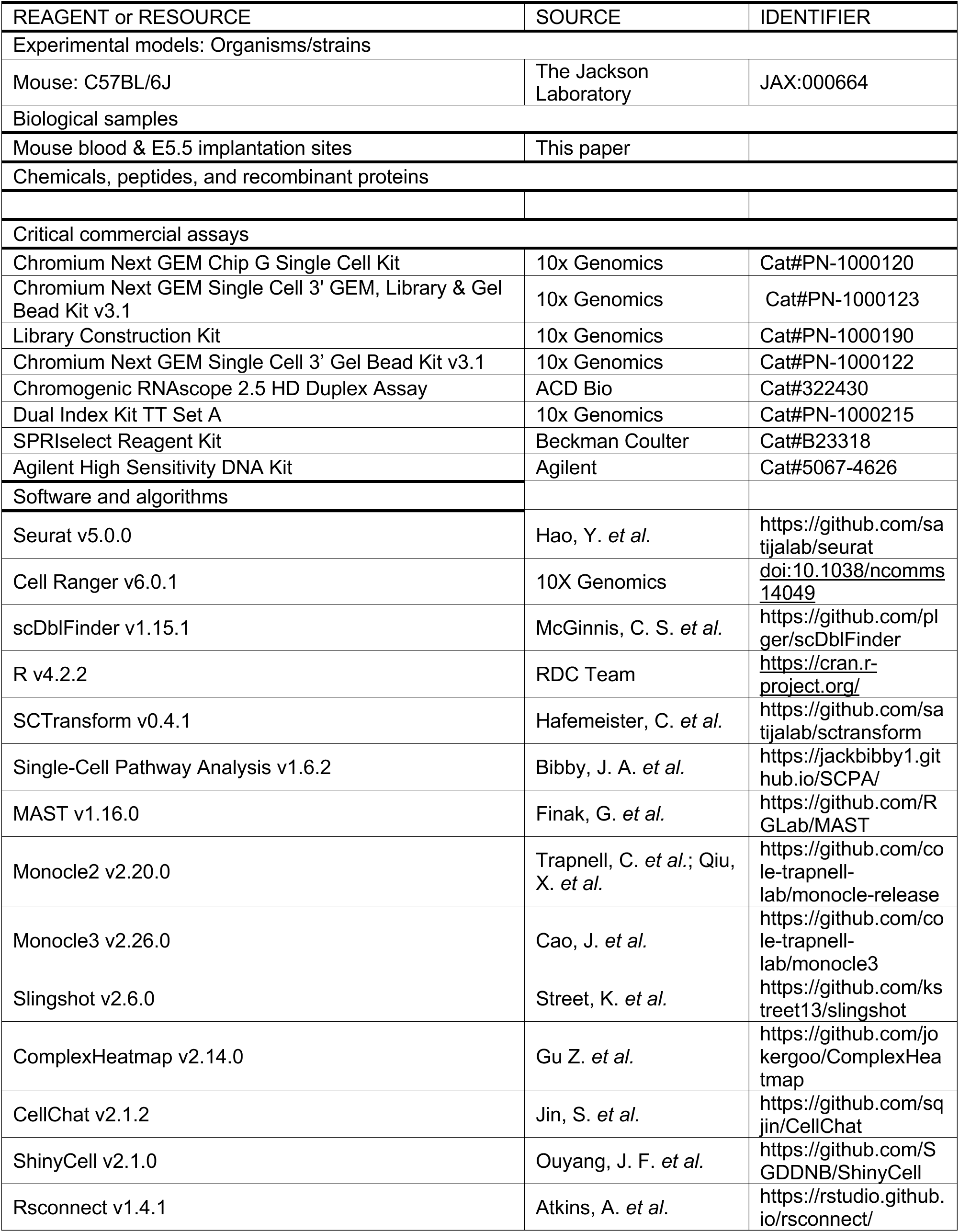
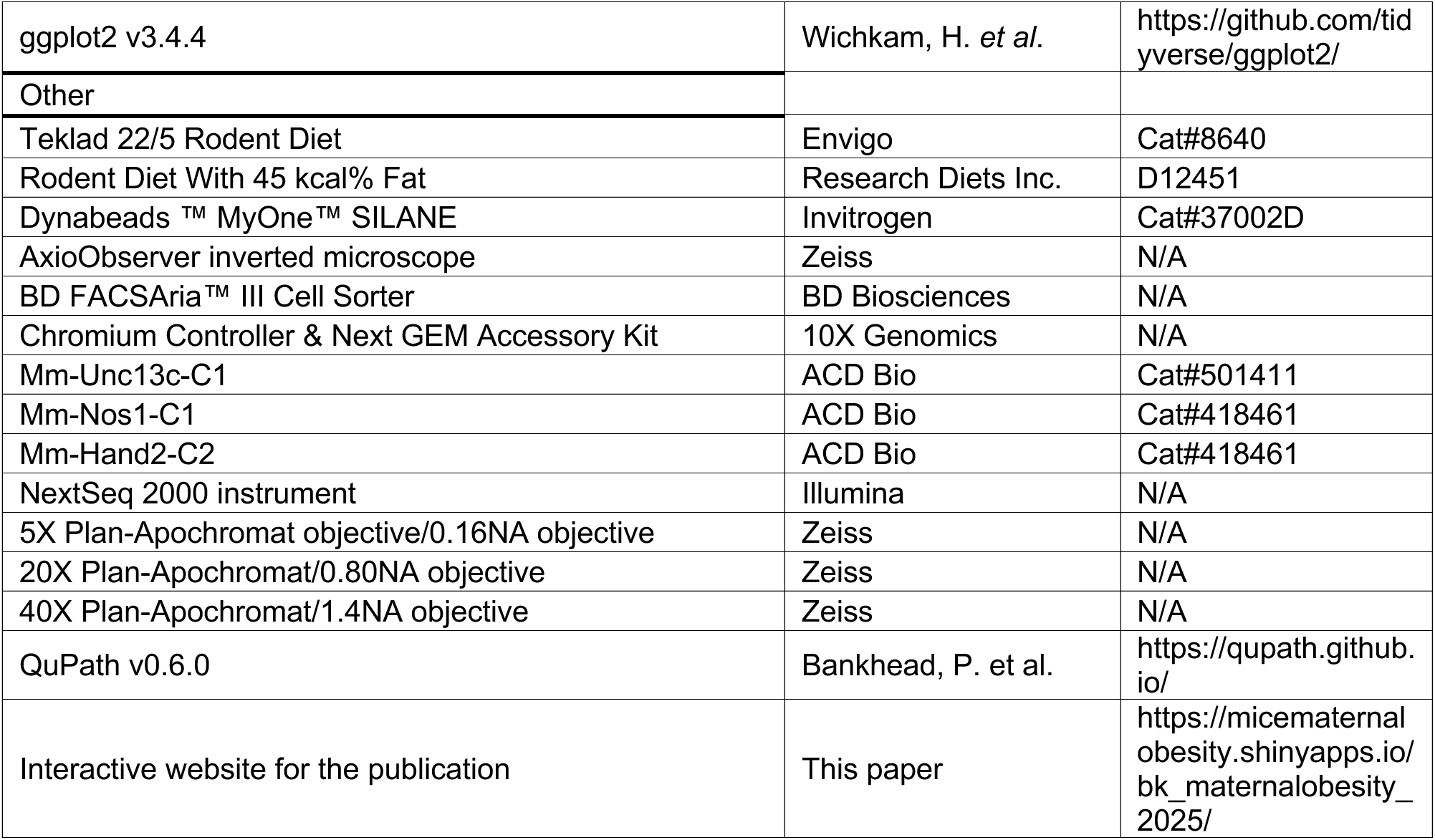

### RESOURCE AVAILABILITY

#### Lead Contact

Alexander G Beristain takes responsibility for the lead contact role. All requests for additional information pertaining to the content of, or materials utilized in, this manuscript should be directed to, and will be addressed by, the lead contact Alexander G. Beristain

#### Materials Availability

This study did not generate new unique reagents. C57BL/6J mice used in this study were purchased from the Jackson Laboratories (Bar Harbor, Maine, USA).

#### Data and Code Availability

- Single-cell RNA-seq data generated in this paper have been deposited at GEO and are publicly available as of the date of publication. The accession numbers for this newly synthesized dataset are listed in the key resources table. Processed sequencing data in the form of Seurat objects are available at: h{ps://micematernalobesity.shinyapps.io/bk_maternalobesity_2025/
- This paper does not report original code. The computational pipelines used for all data processing and bioinformatic analyses in this paper has been deposited at Zenodo and is publicly available as of the date of publication. DOIs are listed in the key resources table.
- Any additional information required to reanalyze the data reported in this paper is available from the Lead Contact upon request.

### Animal Model

#### Ethical statement

All animal procedures for this study were approved by the McMaster University Animal Research Ethics Board (Animal Utilization Protocol 20-07-27) and conducted in accordance with the guidelines of the Canadian Council on Animal Care.

#### Experimental animals

Eight-week-old C57BL/6J female mice (Jackson Laboratories, strain 000664) were randomly assigned to receive either a control rodent chow (Chow; 17% fat, 54% carbohydrate, 29% protein (kCal/g), n = 5; Envigo 8640) or a high fat, high sucrose diet (HFHS; 45% fat, 17% sucrose, 18% carbohydrate, 20% protein (kCal/g), n = 5; Research Diets D12451) *ad libitum* for a minimum of 12 weeks to induce preconception excess adiposity as previously described^32,72,76^. Total fat mass and body weight were measured at baseline and following 10-weeks of diet onset using a whole-body composition analyzer (Minispec LF90-II, Bruker) used to calculate total adiposity relative to bodyweight. Following the dietary intervention, females were time-mated with Chow-fed C57BL/6J males to generate pregnancies. The presence of a copulatory plug in the morning following timed mating was designated embryonic day 0.5 (E0.5), and dams were singly housed until experimental endpoint at embryonic day (E)5.5. At E5.5, dams were euthanized by cervical dislocation and tissues were collected for downstream analysis.

### Tissue collection and dissociation of uterine horns

#### Tissue Collection

Immediately following euthanasia, laparotomy was performed, reproductive tissues were collected, and uterine horns were dissected in Dulbecco’s phosphate-buffered saline (D-PBS, pH 7.4; Wisent Biosciences). The uterus was freed by making incisions at each utero-tubal junction and the cervix. One uterine horn per dam was fixed in 4% w/v paraformaldehyde in D-PBS for 24 hours at room temperature, and the remaining uterine horn was processed to generate single-cell suspensions.

#### Histological confirmation of pregnancy

Fixed uterine horns were processed for paraffin embedding and sectioned at a thickness of 5 µm. The sections were stained with hematoxylin and eosin and examined under brightfield illumination at 20× magnification using a Nikon NiE Eclipse Microscope. Samples displaying focal distension of the uterine wall, lack of a continuous uterine lumen, and cytological changes consistent with decidualization of endometrial stromal cells were used to confirm pregnancy.

#### Tissue dissociation and generation of single cell suspensions

One uterine horn per pregnancy was collected to generate single cell suspensions for transcriptomic profiling using single-cell RNA sequencing (scRNAseq). Tissues were rinsed in Penicillin and Streptomycin supplemented RPMI 1640 basal media (Wisent Biosciences) and minced with scissors. Minced tissue was transferred to a volume of 2.5 ml of tissue digestion media (50 μg/mL Liberase TM (0.28 Wünsch units) and 50 μg/ml DNase I in RPMI). Tissues were digested at 37°C for 30 minutes and filtered through a 70-μm nylon cell sieve into 1mL of FBS. Suspensions were topped up to a volume of 20 mL with RPMI (5% v/v FBS) and centrifuged at 350×g for 10 minutes. Cell pellets were washed and resuspended in 500 µl of D-PBS, and live cell densities in each prep were determined by manual counting with a Neubauer hemocytometer and Trypan Blue-stained aliquots (0.4%; Gibco). Suspensions were pelleted and resuspended in FBS at a density of 2 × 10^6^ live cells/ml. An equal volume of FBS containing 20% dimethyl sulfoxide (DMSO; Millipore Sigma, D2650) was added to cell suspensions as cryoprotectant, for a final suspension of 1 × 10^6^ cells/ml in FBS + 10% DMSO. Samples were cryopreserved at −80°C, then transferred to long-term storage at −150°C until library preparation.

### Generation of scRNAseq library

#### Sample preparation

Cryopreserved cell suspensions from confirmed Chow and HFHS pregnancies (n = 5/group) were rapidly thawed and freezing media removed by washing with RMPI media without glutamine and HEPES (Gibco) supplemented with 10% FBS and 1% Pen/Strep. Cells were pelleted at 400g and resuspended in 500 µl of cell sorting buffer (2% FBS in HBSS 1X Without calcium and magnesium) supplemented with 7-Aminoactinomycin D (7-AAD, 1:40; eBioscience) for 20 minutes to stain dead cells and filtered using a 40 µm FlowMi cell strainer (Bel-Art). Viable (7-AAD negative) cells were isolated using a BD FACS Aria cell sorter using a 100μm nozzle 20 PSI thresholds. Cells were collected in 0.04% UltraPure BSA (Invitrogen) in PBS. Collected cells were used immediately for preparation of single-cell RNA libraries using the Chromium Single Cell 3’ v3.1 kit (10x Genomics) according to the manufacturer’s protocol (CG000315 Rev C) and sequenced on an Illumina NextSeq2000 platform targeting ∼60,000 reads per cell. Sequencing reads were aligned to *mus musculus* reference genome mm10-2.1.0 using cellranger v6.0.1. A total of 97,414 cells were captured for sequencing (Chow n = 49,891 and HFHS n = 47,523) at an average depth of ∼60,000 sequencing reads/cell.

### scRNAseq data analysis

#### Data pre-processing & quality control

Cells were pre-processed using the Seurat R package^37^ (v5.0.0). To exclude low-quality cells, cells with fewer than 1,000 or more than 7,500 detected genes, as well as cells with greater than 10% mitochondrial transcript content were removed. Doublet detection was performed on individual samples using the “scDblFinder” package^77^ (v1.15.1), and predicted doublets were excluded from subsequent analyses. Remaining cells were normalized and scored for cell cycle phase based on canonical G2/M and S phase gene sets. To identify genes exhibiting high biological variability, the “FindVariableFeatures” function was applied using the “vst” selection method, prioritizing the top 2,000 highly variable genes per sample. Data were scaled to center expression values and regress out unwanted sources of variation before integration. Samples were subsequently merged, split by dietary condition, and integrated using the “SCTransform”^78^ v2 workflow. During integration, the Pearson residuals obtained from the default regularized negative binomial model were re-computed and scaled to remove latent variation resulting from the difference between the G2/M and S phase cell cycle scores. After integration, data layers generated through SCTransform were combined into a single layer using the “JoinLayers” function.

#### Cell clustering and cell type annotation

- Integrated object: The top 50 principal components were used for dimensionality reduction and visualization using the Uniform Manifold Approximation and Projection (UMAP) algorithm. Clustering of the single cells was performed at a resolution of 0.24. Cell type annotation was completed by the expression of canonical marker genes, in combination with differentially expressed genes identified using the “FindAllMarkers” function. Marker genes were defined as those exhibiting a log2 fold-change (log2FC) greater 0.1 and an adjusted p-value below 0.01 within a given cluster. If multiple clusters were determined to be same cell type, sub-index is added to that cell type for each different original Seurat cluster. Endometrial stromal cells were determined by expression of *Hoxa10* and *Hoxa11*, immune cells were identified by CD45 (*Ptprc*) expression. Cells expressing *Pecam1* were annotated as endothelial cells, *Krt18* expressing cells were identified as epithelial cells, and *Acta2* and *Myh11* expression was used to determine smooth muscle cells. UMAP plots were generated for integrated object, Chow and HFHS subsets.
- Hand2^+^ ESC subset: Endometrial stromal cell clusters (ESCs) that were also expressing Hand2 were subset from the integrated object and re-clustered in Seurat at a resolution of 0.23 using 30 principal components. Markers were identified using the same strategy as the integrated object. Undifferentiated endometrial stromal cells were annotated based on the expression of *Vim, Snai2, Sfrp4, Ig\p5*. Primary decidual zone (PDZ) clusters were identified by the expression of *H19, Dio3, Krt19, Bmp2*, while secondary decidual zone (SDZ) clusters were defined by the expression of *Scara5, Ig\p3, Dio2, Rb1, Foxp2*. Marker genes were defined by Seurat are those exhibiting log2FC greater than 0.1 and an adjusted p-value below 0.01 within a given cluster.

#### Pseudo-bulk Pearson’s correlation coefficient analysis

Expression matrices from the integrated dataset were averaged by dietary condition and log-transformed to create ‘‘pseudo-bulk’’ gene expression profiles using “AverageExpression” function. Pearson correlation coefficients between pseudo-bulk samples were calculated using the “cor” function from the stats R package (v4.2.2).

#### Differential gene expression

Differential gene expression (DGE) analyses were performed using the “FindMarkers” function in Seurat, implemented with the MAST statistical framework^79^. To maximize transcriptome-wide detection sensitivity, parameter thresholds were set to include all genes regardless of their expression frequency by specifying min.pct = - Inf and min.diff.pct = - Inf. Genes with adjusted p-value below 0.05 and exhibiting a log2FC greater than 0.2 between groups were reported in Figure 2B and Figure 3B. Differentially expressed genes (DEGs) with log2FC greater than 0.5 were plotted as a scatterplot using ggplot2^80^ (v3.4.4). The default RNA assay was used for all analyses.

#### Pathway analysis

Pathway enrichment analysis was performed using a modified implementation of the Single-Cell Pathway Analysis^81^ (v1.6.2) package to evaluate the impact of maternal diet on cellular signaling pathways at single-cell resolution. To increase statistical power and the number of cells contributing to pathway comparisons, the “compare_seurat” function was adapted to allow a downsampling threshold of 3,000 cells per group (default = 500). Gene signatures corresponding to Hallmark pathways were obtained from the msigdbr mouse datasets^82,83^, and formatted for SCPA using the “format_pathways” function. SCPA comparisons were performed across Hand2^+^ ESCs, Hand2^-^ ESCs, natural killer (NK) cells, and macrophages. Pathways enriched in either group with a fold change (FC) > 0.5 and adjusted p-value < 0.01 were visualized using ggplot2^80^ (v3.4.4). To compare pathway-level trends across populations, FC values for each pathway were extracted from the SCPA outputs of Hand2^+^ ESCs, Hand2^-^ ESCs, NK, and macrophage, and a matrix containing combined matrix of enrichment scores was generated. Significant pathways, defined as those meeting FC and adjusted p-value thresholds in at least one population, were used to annotate heatmap rows. Heatmap was generated using the ComplexHeatmap R package^84^ (v2.14.0), displaying FC values across the four cell populations. Unsupervised k-means clustering was applied to both pathways and populations to visualize shared or distinct pathway activity patterns. For Hand2^+^ ESCs, pathway enrichment was assessed using re-clustered object across dietary conditions considering each stromal cluster by defining group1 as diet and group 2 as stromal cell subtypes, whereas for other populations enrichment was compared across dietary condition across the entire cluster.

SCPA was also used to compare pathway activity in distinct Hand2^+^ ESCs sub-types after re-clustering (UnStr, PDZ1, PDZ2, SDZ1, and SDZ2). Pathways enriched in either group with a FC greater than 0.5 and adjusted p-value less than 0.01 were plotted using ggplot2^80^ (v3.4.4).

#### Pseudotime Analysis

The Monocle2 R package^85^ (v2.26.0), Monocle3 R package^86^ (v1.3.1), and Slingshot R package^58^ (v2.6.0) were used to explore the differentiation of undifferentiated ESCs to PDZ and SDZ endpoints in Hand2^+^ ESCs.

1. *Monocle2:* Re-clustered Hand2^+^ ESC v5 RNA assay was first converted to a Monocle2-compatible v3 RNA assay following the standard Seurat RNA5-to-RNA3 pipeline and Seurat object was subsequently converted to CellDataSet object using “as.CellDataSet” function. Cells were then imported into monocle2 to perform semi-supervised single-cell ordering in pseudotime using Sfrp4, IgÖp5, Sparc, Fst, Mfap4, Ogn, Fos to denote cells at the origin. Progression towards PDZ was characterized by Dio3, Id3, and Ran whereas progression towards SDZ was characterized by Scara5, Apoe, and Inmt. Branched expression analysis modelling (BEAM) was applied to assess gene expression dynamics along bifurcating pseudotime trajectories progressing towards PDZ or SDZ maturation. Trajectory graphs were constructed by first reducing dimensions using “reduceDimension” function, followed by “orderCells” function to order cells in pseudotime designating UnStr as the origin based on the known transcriptomic markers. Trajectory ordering and structures were visualized using “plot_cell_trajectory” function. For comparative analyses, re-clustered Hand2^+^ ESCs were further subset based on dietary condition (Chow and HFHS) in order to generate diet-specific single-cell pseudotime ordering, and independent trajectories were constructed using the same genes and gene expression thresholds for diet-specific comparison. In order to test each gene for differential expression analysis as a function of pseudotime, differential gene test was performed with the object containing both Chow and HFHS cells using “differentialGeneTest” function by combining monocle2 pseudotime and diet information as covariates, and select genes were plo{ed using “plot_genes_branched_heatmap” function for Chow and HFHS objects along the pseudotime.
2. *Monocle3:* For trajectory modeling with Monocle3, re-clustered Hand2^+^ ESC expression matrices were imported, clustered, and used to create the principal graph with the “learn_graph” function. Cells were ordered in pseudotime using “order_cells” function with the trajectory root designated based on the root cell ID identified in the corresponding Monocle 2 analysis. To visualize lineage trajectories in three-dimensional space, “plot_cells_3d” function was used and two distinct, branching trajectories corresponding to PDZ and SDZ maturation was shown.
3. *Slingshot:* To perform trajectory modeling using Slingshot, re-clustered Hand2^+^ ESC object was converted to a SingleCellExperiment object using “as.SingleCellExperiment” function. Lineages were constructed in UMAP space using the “slingshot” function, with the unStr cluster specified as the starting point and PDZ2 and SDZ2 clusters as the endpoints. A stretch factor of 0 and default lineage extension parameters were applied. Pseudotime values along each lineage were calculated and visualized by projecting cells onto the UMAP embedding, with colors representing pseudotime values for each cell. Lineage curves were overlaid using the lines function from Slingshot to illustrate differentiation pathways, and pseudotime_1 and pseudotime_2 values were plo{ed in UMAP space. For comparative analyses, re-clustered Hand2^+^ ESCs were further subset based on their cluster identity. The UnStr, PDZ1 and PDZ2 were used to generate PDZ-specific single-cell pseudotime ordering while UnStr, SDZ1, and SDZ2 were subset to model SDZ-specific pseudotime ordering. Independent trajectories were constructed using the same parameters used for the initial integrated analysis. To assess the effect of maternal diet on ESC differentiation, pseudotime distributions along the PDZ and SDZ trajectories were compared between Maternal Chow and HFHS groups using a density plot generated with ggplot2^80^ (v3.4.4), and statistical significance was assessed using the Kolmogorov-Smirnov Test using “ks.test” function from stats R package (v4.2.2).

#### Cell-cell interaction analysis

Cell-cell communication was analyzed using CellChat R package^59^ (v2.1.2) to infer ligand-receptor signaling networks in early post-implantation mice uterus following the author’s pipeline. Object containing between Hand2^+^ ESC subsets, NK and macrophages was subset based on diet, and Chow and HFHS independently processed using CellChat to create Chow and HFHS data objects containing inferred interactions. Normalized gene expression matrix and cell-level metadata were provided as input to CellChat, with group identities defined by stromal subset annotation. The default ligand-receptor interaction database for mouse “CellChatDB.mouse” was used to infer potential intercellular communications. Overexpressed genes and ligand-receptor interactions were identified using the “subsetData”, “identifyOverExpressedGenes”, and “identifyOverExpressedInteractions” functions. Communication probabilities were computed using the trimean method “computeCommunProb” and filtered for interactions using “filterCommunication” containing at least 10 cells per interacting group. Inferred communications were aggregated at both the ligand-receptor pair and signaling pathway levels using “computeCommunProbPathway” and “aggregateNet”. Network centrality analysis was performed to assess signaling roles of each cell group, and significant communication networks were visualized using circle plots “netVisual_aggregate”.

Differential communication analyses between dietary conditions were conducted by merging CellChat objects with “mergeCellChat”, followed by comparative network visualization of top 50% interactions using “netVisual_diffInteraction” and RankNet analysis “rankNet” to quantify differential signaling contributions. Communication patterns were identified separately for outgoing and incoming signals using “identifyCommunicationPatterns” based on unsupervised clustering of signaling probabilities. Heatmap was generated using “netVisual_heatmap” function to show differential number of interactions and interaction strength in the cell-cell communication network between two dietary conditions. For visualization of specific pathways such as Desmosterol and Galectin, both the interaction networks were plotted using “netVisual_aggregate”. Ligand and receptor expressions of these two pathways were plotted after identifying significant ligand-receptor pairs and extracting related genes using “extractEnrichedLR” function, and plotting them using Seurat “VlnPlot” function for Desmosterol and Galectin signaling pathways, respectively. Increased or decreased significant ligand-receptor pairs from one cell type to other cell type were plotted using “netVisual_bubble” function by specifying the source cluster and target clusters.

### RNA Scope in situ hybridization

Tissues that were fixed and paraffin embedded for histological confirmation of pregnancy were sectioned at 5 μm and mounted onto glass slides. The chromogenic RNAscope 2.5 HD Duplex Assay (ACD Bio) was performed according to manufacturer’s instructions. The Mm-Unc13-C1 (ACD bio), Mm-Nos1-C1 (ACD bio), and Mm-Hand2-C2 (ACD bio) probes were used. Slides were imaged with an AxioObserver inverted microscope (Zeiss) using the 5X, 20X, and 40X Plan-Apochromat/0.8NA objective (Zeiss). Quantification of *in situ* RNAScope signals were performed using Qupath software ^87^

### Interactive App Generation

The ShinyCell R package^88^ (v2.1.0) was used to assemble the “Mice Maternal Obesity” app and the rsconnect R package (v1.4.1) was used to publish the app on the shinyapps.io server for public access.

## QUANTIFICATION AND STATISTICAL ANALYSIS

### Statistical analysis

All statistical analyses in this study were performed in R (v.4.4.2; https://www.R-project.org). One dam or pregnancy was considered a biological replicate. The normality of all data sets was determined using the D’Agostino-Pearson K2 test. Pairwise comparisons were made using univariate linear models with maternal diet as the dependent variable, or Mann-Whitney U-test where the data were discrete values or non-normally distributed. For time series experiments or those with repeated/replicate measures, linear mixed effects models were constructed using the lmerTest package ^89^ (version 3.1-3, RRID SCR_015656) and analyzed using two-way ANOVA with Type 3 sum of squares and Kenward-Roger calculated denominator degrees of freedom using the Anova function in the car package (version 3.1-3, RRID SCR_022137). For these analyses, maternal diet and time were set as fixed effects, and dam as a random effect for nesting repeated measurements. Where significant main effects or interactions were present, pairwise post hoc comparisons were made between the estimated marginal means using Tukey’s HSD method with the emmeans package ^90^ (v.1.8.5, RRID SCR_018734). For all analyses, p < 0.05 was used as a nominal threshold for determining statistical significance. Please refer to the relevant scRNAseq analysis methods for statistical analyses applied to single-cell datasets.

## EXCEL TABLE TITLES AND LEGENDS

Table S1: Metadata, related to Methods, Figure 1 and Supplemental Figure 1

Table S2: Full list of gene markers, related to Supplemental Figure 1

Table S3: List of DEGs in E5.5 endometrial cell types as a consequence of obesogenic diet, related to Figure 2.

Table S4: List of enriched gene pathways in endometrial stromal cells, uNK, and Mϕ as a result of obesogenic diet, related to Figure 2.

Table S5: Gene markers of *Hand2*^+^ endometrial stromal cell subtypes, related to Methods, Figure 3, and Supplemental Figure 3.

Table S6: List of DEGs in *Hand2*^+^ endometrial stromal cell types as a consequence of obesogenic diet, related to Figure 3 and Supplemental Figure 3.

Table S7: List of enriched gene pathways in Hand2+ endometrial stromal cells as a consequence of obesogenic diet, related to Figure 3 and Supplemental Figure 3.

Table S8: CellChat rankNet signaling contribution results, related to Figure 5.

